# Biochemical insight into novel Rab-GEF activity of the mammalian TRAPPIII complex

**DOI:** 10.1101/2021.06.01.446621

**Authors:** Noah J Harris, Meredith L Jenkins, Udit Dalwadi, Kaelin D Fleming, Sung-Eun Nam, Matthew AH Parsons, Calvin K Yip, John E Burke

**Affiliations:** Department of Biochemistry and Microbiology, University of Victoria, Victoria, British Columbia, Canada V8W 2Y2; Life sciences Institute, Department of Biochemistry and Molecular Biology, The University of British Columbia, Vancouver, British Columbia V6T 1Z3, Canada

**Keywords:** Rab GTPases, Rab1, Rab43, TRAPP, TRAPPII, TRAPPIII, electron microscopy, HDX-MS, hydrogen deuterium exchange

## Abstract

Transport Protein Particle complexes (TRAPP) are evolutionarily conserved regulators of membrane trafficking, with this mediated by their guanine nucleotide exchange factor (GEF) activity towards Rab GTPases. In metazoans evidence suggests that two different TRAPP complexes exist, TRAPPII and TRAPPIII. These two complexes share a common core of subunits, with complex specific subunits (TRAPPC9 and TRAPPC10 in TRAPPII and TRAPPC8, TRAPPC11, TRAPPC12, TRAPPC13 in TRAPPIII). TRAPPII and TRAPPIII have distinct specificity for GEF activity towards Rabs, with TRAPPIII acting on Rab1, and TRAPPII acting on Rab1 and Rab11. The molecular basis for how these complex specific subunits alter GEF activity towards Rab GTPases is unknown. Here we have used a combination of biochemical assays, hydrogen deuterium exchange mass spectrometry (HDX-MS) and electron microscopy to examine the regulation of TRAPPII and TRAPPIIII complexes in solution and on membranes. GEF assays revealed that the TRAPPIII has GEF activity against Rab1 and Rab43, with no detectable activity against the other 18 Rabs tested. The TRAPPIII complex had significant differences in protein dynamics at the Rab binding site compared to TRAPPII, potentially indicating an important role of accessory subunits in altering the active site of TRAPP complexes. Both the TRAPPII and TRAPPIII complexes had enhanced GEF activity on lipid membranes, with HDX-MS revealing numerous conformational changes that accompany membrane association. HDX-MS also identified a membrane binding site in TRAPPC8. Collectively, our results provide insight into the functions of TRAPP complexes and how they can achieve Rab specificity.

## Introduction

Membrane trafficking is an essential process in all eukaryotic cells and requires tightly coordinated transport of cellular material to distinct intracellular organelles. Numerous different proteins are involved in regulating this pathway but one of the most important protein families involved are Rab GTPases. Rabs act as molecular switches that cycle between GTP-bound active and GDP-bound inactive states. Rabs recruit downstream effector molecules depending on their nucleotide binding state, including molecular motors, and tethers which control membrane trafficking events. The activation of Rabs is catalyzed by Guanine nucleotide exchange factors (GEFs) [1–3] which mediate the exchange of GDP for GTP. Defining how Rabs are targeted by GEFs is important in understanding their function in membrane trafficking. The Transport Protein Particle (TRAPP) complexes are evolutionarily conserved in all eukaryotic cells, and are potent Rab GEFs playing important roles in secretion, autophagy, and Golgi trafficking [4–9].

Metazoans are proposed to form two different TRAPP complexes: TRAPPII and TRAPPIII. TRAPPII can activate Rab1, but it has been proposed that its main role is to activate Rab11, therefore playing key roles in secretion from the Golgi [10–13]. TRAPPIII does not have activity against Rab11, and instead is proposed to primarily mediate activation of Rab1, an important regulator in ER-Golgi, intra-Golgi trafficking and autophagy [11,14–16]. The mammalian TRAPPII complex has additional activity on Rab19 and Rab43 [10]. Rab19 and Rab43 are both Golgi-localised Rabs, with Rab43 being involved in mediating GPCR trafficking [17]. Defining the molecular mechanisms that mediate GEF specificity of mammalian TRAPP complexes will be critical in understanding their roles in membrane trafficking.

TRAPPI, TRAPPII and TRAPPIII all share seven conserved subunits that make up the TRAPP “core” (TRAPPC1, TRAPPC2, TRAPPC2L, TRAPPC3A/B, TRAPPC4, TRAPPC5, TRAPPC6A/B) [18]. Mammalian TRAPPII is composed of the core and two additional complex specific subunits (TRAPPC9, TRAPPC10) with mammalian TRAPPIII composed of the core and four additional complex specific subunits (TRAPPC8, TRAPPC11, TRAPPC12, TRAPPC13) [11,19,20]. The importance of TRAPPIII as a Rab1 GEF is highlighted by TRAPPC8 and TRAPPC11 being essential for cell survival and Rab1 activation [11,21]. Highlighting the critical role of TRAPP complexes in human disease is that mutations or deletions in TRAPP specific subunits have been found to be involved in several neurodevelopmental disorders collectively known as “TRAPPopathies” [5,6,9].

Structural studies using X-ray crystallography and electron microscopy have revealed the architecture of the conserved core of the TRAPP complexes and how they associate with Rab GTPases (Fig. 1A) [18,22]. The structure of the yeast TRAPP core complex bound to Ypt1 (Rab1 homolog) revealed the Rab binding interface which is composed of TRAPPC1, TRAPPC3, and TRAPPC4 [18,22]. The cryo-electron microscopy (cryo-EM) structure of the yeast TRAPPIII complex revealed the interactions of the core with TRAPPC8, with an additional interaction of TRAPPC5 with the hyper-variable tail (HVT) of Ypt1 (Rab1 homolog) [23]. The cryo-EM structure of the Drosophila TRAPPIII complex bound to Rab1 shows how the additional TRAPPIII complex specific subunits are arranged in relation to the core, with a novel interaction between TRAPPC8 and Rab1 [24]. Despite these insights into the assembly of TRAPPIII complexes, it is still not understood how the TRAPPIII complex specific subunits change the way the conserved subunits interact with Rab substrates, and how substrate selectivity is controlled.

**Figure 1.**
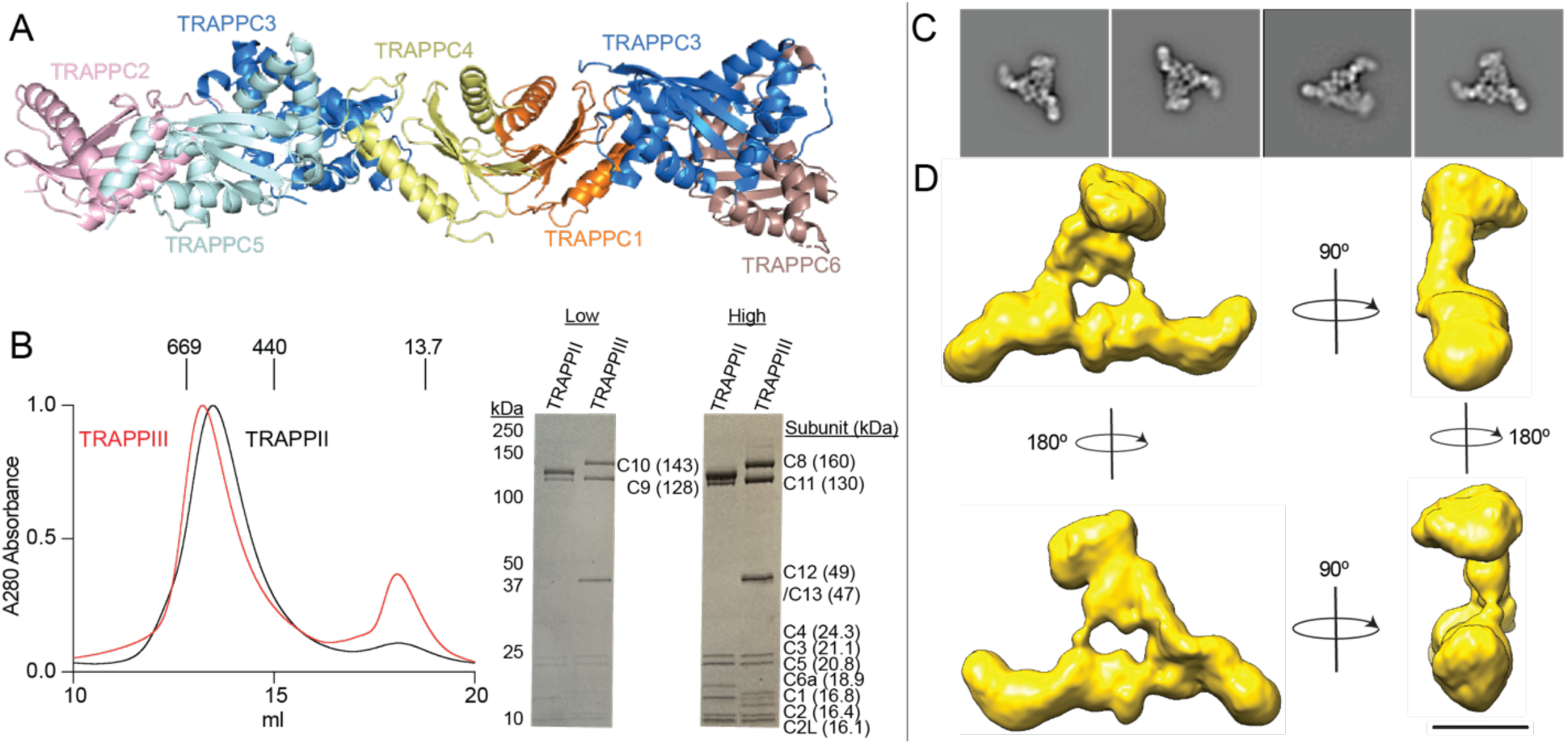
Purification and Architecture of the mammalian TRAPPIII complex. **A.** Model of the mammalian TRAPP conserved subunits. The model was generated through a combination of the following structures (PDB: 2J3T, 2J3W, and 3CUE) Phyre2 was used to generate structures of mammalian TRAPP subunits with no solved crystal structure [26]. **B.** left: Size Exclusion Chromatography (SEC) trace of TRAPPII and TRAPPIII on a Superose 6 gel filtration column with molecular weight markers indicated in kDa. Absorbance is normalized to max absorbance. Right: SDS-PAGE gel of purified mammalian TRAPPII and TRAPPIII complex. Protein composition is shown on the right. High and low labels refer to protein amount loaded (high=3.6 *µ*g and low= 1.4 *µ*g). **C.** Representative 2D negative stain electron microscopy class averages of the TRAPPIII complex. Box edge length is 670 Å. **D.** 3D EM reconstruction of TRAPPIII with different orientations of the complex. Scale bar represents 100 Å.

Here we have carried out biochemical and biophysical analysis of the mammalian TRAPPIII complex using a combination of in-vitro GEF assays, electron microscopy, and hydrogen deuterium exchange mass spectrometry (HDX-MS). We have purified both mammalian TRAPPII (composed of TRAPPC1, TRAPPC2, TRAPPC2L, TRAPPC3, TRAPPC4, TRAPPC5, TRAPPC6A, TRAPPC9, TRAPPC10) and TRAPPIII (composed of TRAPPC1, TRAPPC2, TRAPPC2L, TRAPPC3, TRAPPC4, TRAPPC5, TRAPPC6A, TRAPPC8, TRAPPC11, TRAPPC12, TRAPPC13). Single-particle electron microscopy analysis of the purified TRAPPIII revealed an overall architecture resembling that of TRAPPIII isolated from Drosophila. We characterised TRAPPIII GEF activity against 20 different Rab GTPases both in solution and on membranes. We found TRAPPIII mediated GEF activity for Rab1 and Rab43, but no activity towards other any other Rab GTPases tested. HDX-MS comparing the TRAPPII and TRAPPIII complexes showed extensive differences at the Rab binding site, revealing a potential role of complex specific subunits in reshaping the GEF catalytic site. GEF activity of both TRAPPII and TRAPPIII was enhanced on membranes, with HDX-MS revealing extensive conformational changes upon membrane binding and identifying a conserved membrane binding site in the TRAPPIII specific subunit TRAPPC8. Overall, this work provides unique insight into the architecture and dynamics of TRAPP complexes, and the mechanisms by which their GEF activity is regulated.

## Results

### Purification and architecture of the mammalian TRAPPIII complex

We recombinantly purified TRAPPII and TRAPPIII using a similar approach to our previous work on TRAPPII [10]. TRAPPII and TRAPPIII complexes were generated using the biGBac multi-promoter system [25] in *Sf9* insect cells and protein purification was carried out using HisTrap and StrepTrap affinity columns followed by gel filtration (Fig. 1B). The details outlining the specific plasmids and TRAPP subunit boundaries used can be found in Table S1. TRAPPIII eluted at a size consistent with one copy of all subunits (exception being two copies of TRAPPC3) on size exclusion chromatography. Tandem mass spectrometry (MS/MS) analysis of the purified TRAPPII and TRAPPIII complexes identified peptides covering all expressed subunits.

To investigate the architecture of the mammalian TRAPPIII complex we subjected purified mammalian TRAPPIII to negative stain single particle electron microscopy (EM) analysis (Fig. 1C-D, Fig. S1). Raw images revealed triangular-shaped particles similar to Drosophila TRAPPIII [24] but distinct from the smaller-sized yeast TRAPPIII [23]. Two-dimensional (2D) analysis and 3D reconstruction revealed mammalian TRAPPIII is composed of an elongated rod-like region reminiscent of the TRAPP core (TRAPPC1, TRAPPC2, TRAPPC3A/B, TRAPPC4, TRAPPC5, TRAPPC6A), two “arms” capping the ends of the putative TRAPP core, and a peripheral region extending from the core (Fig. 1D). The ultrastructure of mammalian TRAPPIII is consistent with the cryo-EM structure of Drosophila TRAPPIII which identified a triangular-shaped complex with a similar rod-like TRAPP core that contains TRAPPC8 and TRAPPC11 “arms” on either end of the core with a peripheral TRAPPC12/C13 region extending from this core region. This suggests a highly conserved architecture between TRAPPIII complexes in metazoans.

### Determination of mammalian TRAPPIII Rab Specificity

To examine the specificity of the GEF activity of the mammalian TRAPPIII complex we tested 20 different Rab GTPases. We focused on Rab GTPases that are evolutionarily similar to Rab1 and Rab11, to identify possibly overlooked Rabs. GEF assays were carried out using Rabs preloaded with the fluorescent GDP analog 3-(N-methyl-anthraniloyl)-2-deoxy-GDP (Mant-GDP) and nucleotide exchange was determined as a function of TRAPP concentration. Rab GTPases were designed with a C-terminal his tag which allowed for GEF activity to be measured in the presence of NiNTA containing synthetic membranes.

In solution, the TRAPPIII complex had GEF activity on Rab1 with a catalytic efficiency of ∼1.7×10^3^ M^−1^s^−1^ which is similar to the catalytic efficiency that we previously determined for TRAPPII on Rab1 (2.9×10^3^ M^−1^s^−1^) and consistent with TRAPPIII’s role as a Rab1 GEF [11,14] (Fig. 2C). Similar to TRAPPII, we found TRAPPIII mediated activity towards Rab43. TRAPPIII showed increased catalytic efficiency for Rab43 compared to Rab1 (5.2×10^3^ M^−1^s^−1^ vs 1.7×10^3^ M^−1^s^−1^, respectively) which was a similar trend to what was observed with the mammalian TRAPPII complex (Fig. 2C) [10]. There was no detectable TRAPPIII mediated GEF activity for any other Rab GTPase tested including Rab11a, Rab11b, or Rab19 which are substrates for mammalian TRAPPII (Fig. 2D). This is consistent with TRAPPIII having GEF activity for Rab1 but not Rab11 in both metazoans and yeast [11,27]. The lack of GEF activity for TRAPPIII with Rab19 was striking because Rab43 and Rab19 are very evolutionarily similar to each other, only diverging in vertebrata with TRAPPII being able to activate both Rab19 and Rab43. A conservational alignment of these GTPases is shown in supplemental figure 2. Collectively, our *in-vitro* analysis of TRAPPIII shows that TRAPPIII is a specific GEF for Rab1a and Rab43 in solution and reveals insight into an unexpected difference in GEF activity between TRAPPII and TRAPPIII towards Rab19. There is no clear sequence relationship explaining the selectivity of why TRAPPII, but not TRAPPIII, is active on Rab19/Rab11 (Fig. S2), presenting a need for further high resolution structural studies to help define this.

**Figure 2.**
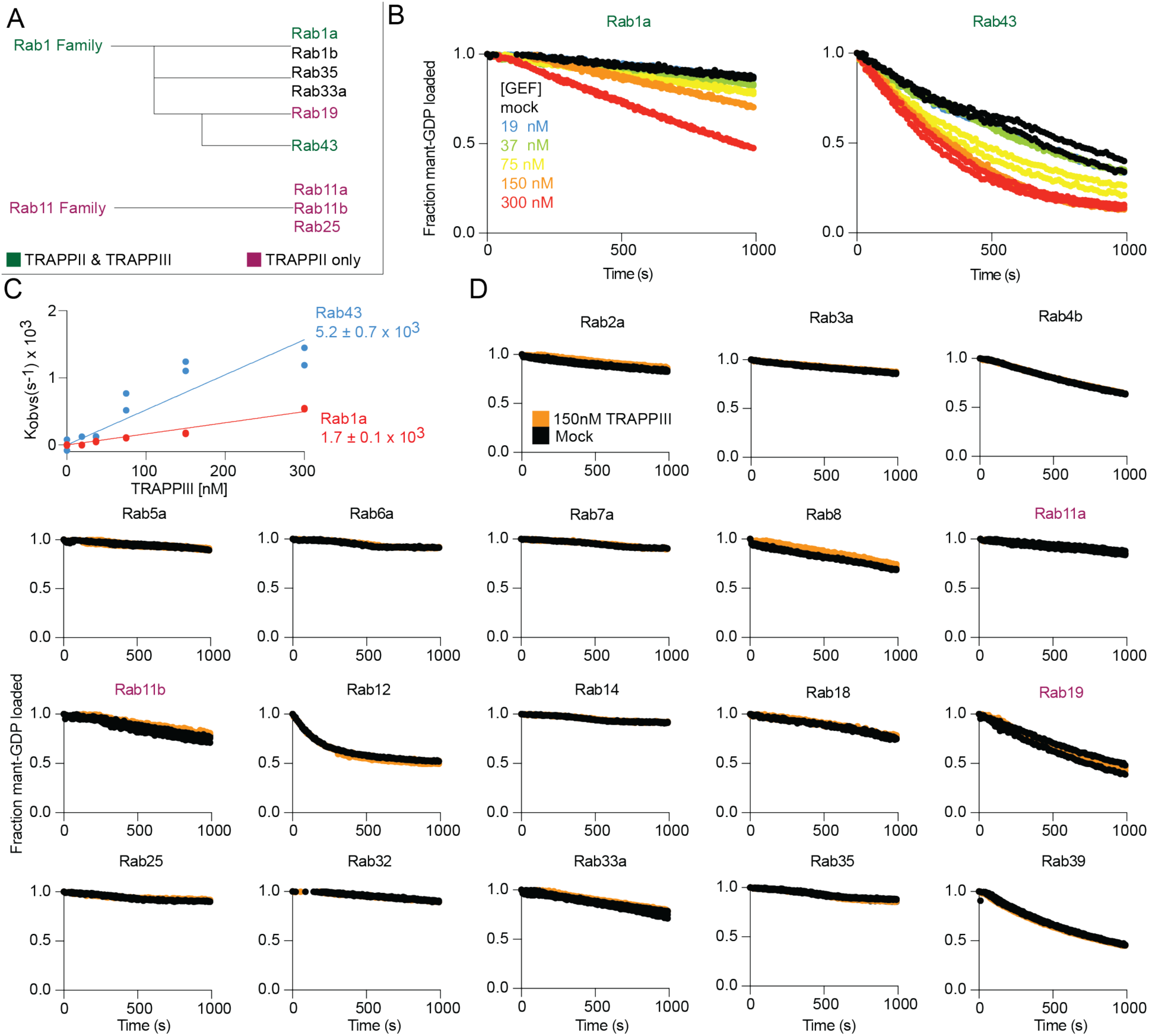
Biochemical analysis of the mammalian TRAPPIII complex against a panel of different Rab GTPases. **A.** A simplified Rab evolutionary schematic (based on [28]). Rabs are colored according to their TRAPPII and TRAPPIII GEF activity. **B.** In-vitro GEF assay of TRAPPIII on Rab1a and Rab43. Nucleotide exchange was monitored by measuring the fluorescent signal during the TRAPPIII (19 nM-300 nM) catalyzed release of Mant-GDP from 4 *µ*M of Rab-His6 in the presence of 100 *µ*M GTPγS. Each concentration was conducted in duplicate (n=2). **C.** Nucleotide exchange rates of Rab1 and Rab43 plotted as a function of TRAPPIII concentration (n=2). **D.** In vitro GEF assay of TRAPPIII on the panel of Rab GTPases. 4 *µ*M Rab was loaded with Mant-GDP in the presence of 150nM TRAPPIII. Each concentration was either conducted in duplicate or triplicate (n=3 for Rab11a, Rab11b, Rab18, Rab19, Rab33, Rab35, n=2 for the others).

### Determining conformational differences upon Rab1 binding to mammalian TRAPPIII

We carried out HDX-MS experiments of TRAPPIII binding to Rab1 to understand the molecular basis for how TRAPPIII binds to and activates Rab1. HDX-MS measures the exchange rate of amide hydrogens in solution, with the rate being primarily determined by stability of secondary structure. It is a powerful tool to measure protein conformational dynamics. Deuterium incorporation in HDX-MS requires the generation of pepsin peptide fragments spanning the entire complex. We obtained peptide maps spanning 69-97% of the entire complex (**Core**: TRAPPC1, TRAPPC2, TRAPPC2L, TRAPPC3, TRAPPC4, TRAPPC5, TRAPPC6A; **TRAPPII specific**: TRAPPC9, TRAPPC10; **TRAPPIII specific**: TRAPPC8, TRAPPC11, TRAPPC12, TRAPPC13), with specific coverage values listed in Table S2-S4. Significant differences in exchange between conditions were defined as differences at any timepoint fitting the following three criteria (>5%, >0.5 Da, and two tailed T-test p value <0.01).

HDX-MS experiments were performed in the presence of EDTA to generate a nucleotide free stabilised Rab-GEF complex. Experiments were carried out under three conditions: EDTA treated Rab, TRAPPIII alone, and TRAPPIII bound to Rab1. We observed decreased exchange upon formation of the Rab-TRAPPIII complex in TRAPPC4 (5-19, 112-136, 180-191) and Rab1a (49-79) (Fig. 3B+C). There were multiple regions with increased exchange in Rab1a (12-40, 91-108, 116-165), likely driven by GEF mediated nucleotide loss. The full HDX-MS data for all subunits and Rab1a is outlined in the source data. Decreases in exchange in TRAPPC4 (180-191) mapped onto the putative Rab binding site consistent with this peptide being protected when Rab was incubated with TRAPPII [10]. Decreases in exchange in Rab1 were observed in the interswitch and switchII regions upon binding the complex (49-79) (Fig. 3B+C). Rab1 was also destabilized in the nucleotide binding pocket as would be expected with the loss of nucleotide upon binding its GEF. No other significant changes were observed in the TRAPPIII complex upon Rab binding. These results suggest that both mammalian TRAPPII and TRAPPIII complexes are binding Rabs at a similar interface.

**Figure 3.**
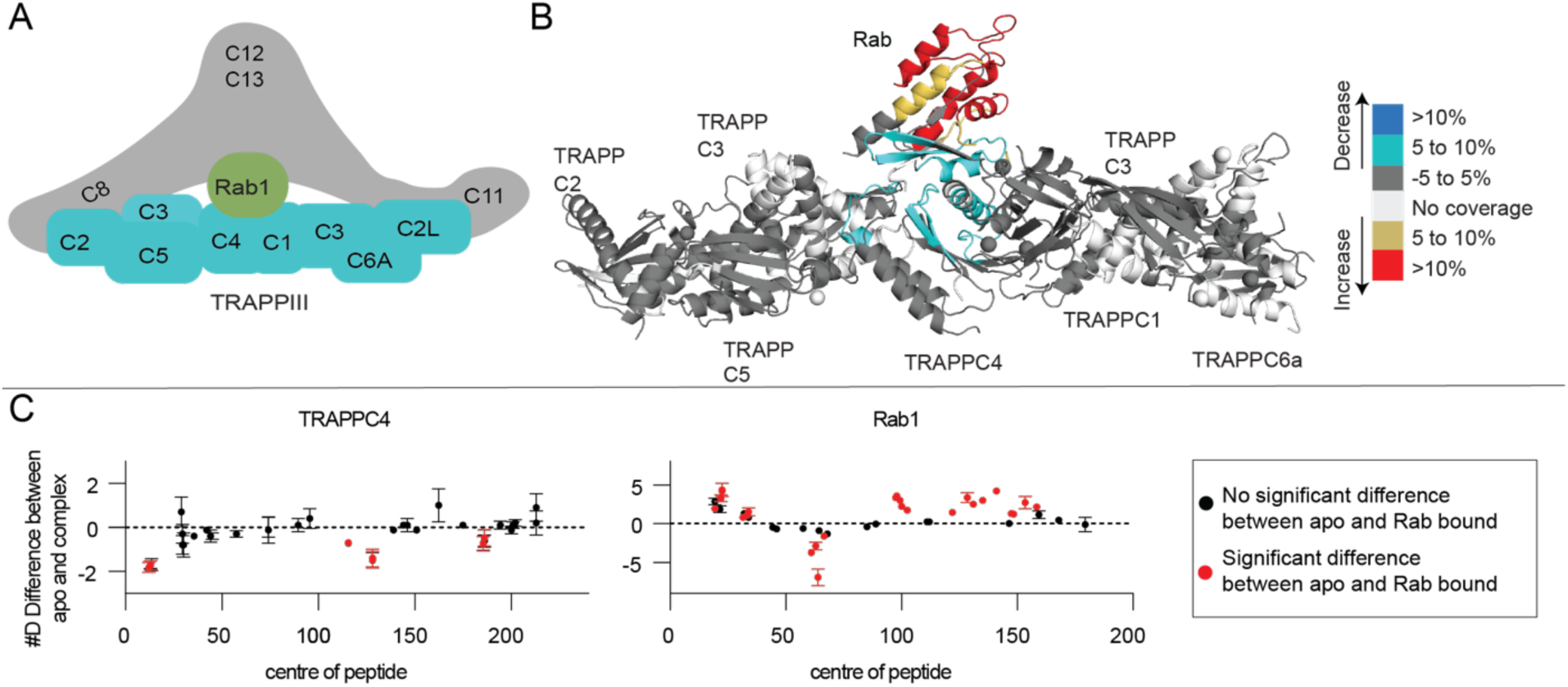
Defining the Rab binding site in the mammalian TRAPPIII complex. **A.** Cartoon schematic highlighting TRAPPIII with bound Rab GTPase. **B.** Significant differences in HDX observed across all time points when Rab1 was incubated with TRAPPIII are mapped onto the predicted model of the TRAPP core. Regions with differences are coloured according to the legend. **C.** The number of deuteron differences for all analysed peptides over the entire deuterium exchange time course for TRAPPIII in the presence of Rab1 (n=3). Only subunits with significant differences (defined as >5%, >0.5 Da, two tailed T-test p value <0.01), are shown. Every point represents the central residue of an individual peptide with significant differences being indicated by points colored red.

### Dynamic differences exist at the Rab binding site of the core subunits in the mammalian TRAPPII and TRAPPIII complexes

We carried out hydrogen deuterium exchange mass spectrometry (HDX-MS) experiments comparing TRAPPII vs TRAPPIII to understand any conformational differences that occur in the core between the two complexes. H/D exchange differences between TRAPPII and TRAPPIII can be analyzed only in the shared core components. Multiple different subunits had significant differences in deuterium exchange. TRAPPIII was more protected than TRAPPII in TRAPPC2L (72-80, 100-135), and TRAPPC4 (137-155, 170-204) (Fig 4B). TRAPPII was more protected than TRAPPIII in TRAPPC2 (47-53, 59-70), TRAPPC2L (17-25, 37-66, 81-99), TRAPPC4 (65-83) and TRAPPC5 (58-69, 180-188) (Fig 4B). The full HDX-MS data for all subunits is summarized in the source data. The TRAPPIII complex was more protected than TRAPPII at the canonical Rab interface in TRAPPC4 (170-204). This region in TRAPPC4 would likely be in contact with the N and C termini of the Rab substrate (Fig 4C). Intriguingly, this is in the same region that was protected in TRAPPC4 (181-191) with Rab1a (Fig 3C). The large protections observed in the TRAPPC2L subunit are likely due to interactions with TRAPPIII specific subunits which is consistent with cryo-EM data showing TRAPPC2L being at the interface with TRAPPC11 in the Drosophila TRAPPIII complex [24]. Regions of TRAPPC2 were more protected in the TRAPPII complex, which is unexpected since this is the putative binding interface for the TRAPPIII specific subunit TRAPPC8. Together these data suggest that TRAPPII and TRAPPIII have dynamic differences in their core subunits that are involved in Rab binding, which may be playing a role in mediating Rab specificity.

**Figure 4.**
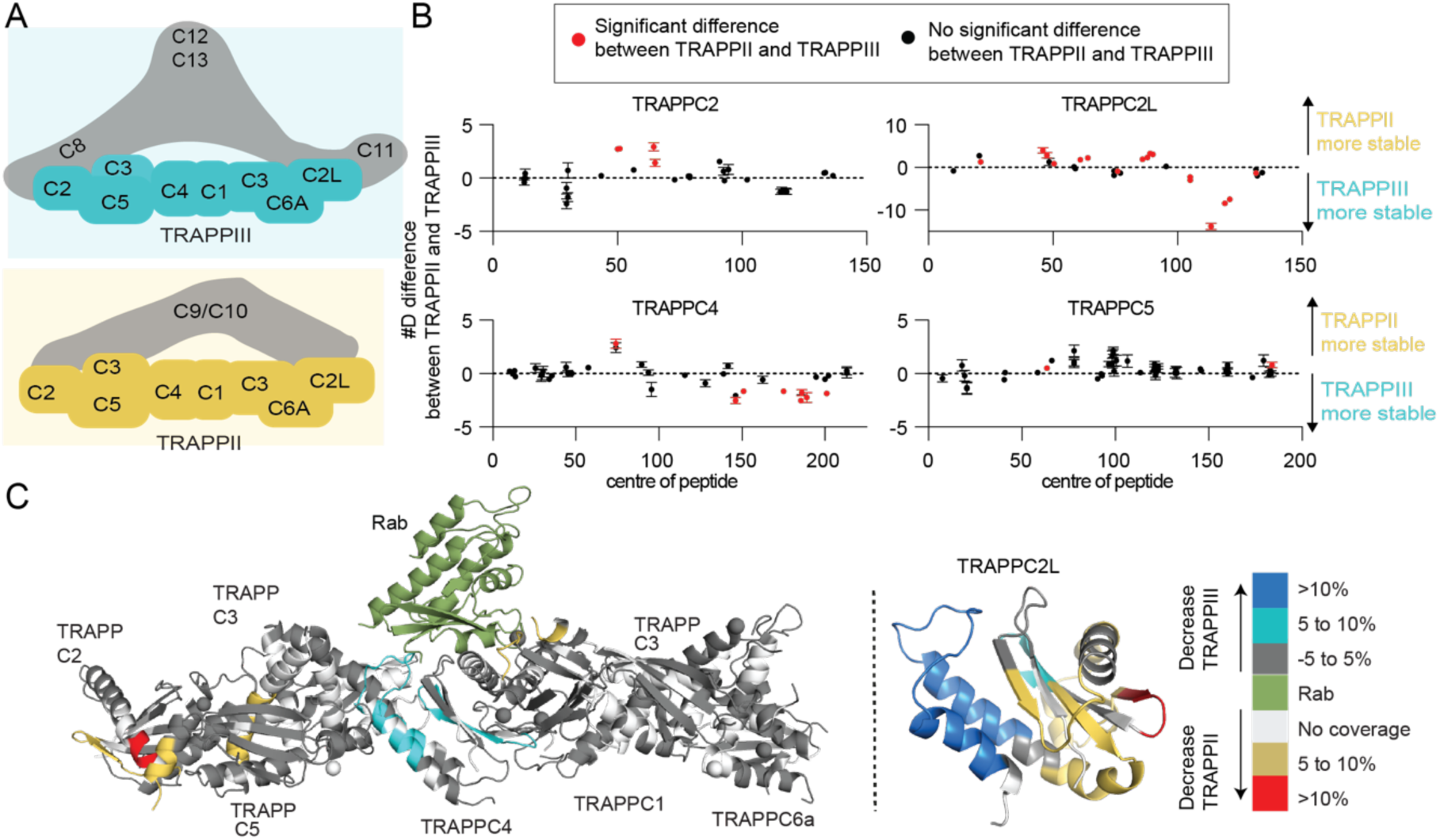
Comparative HDX-MS reveals dynamic differences within the core of TRAPPIII compared to TRAPPII. **A.** Cartoon schematic highlighting protections or destabilisations in TRAPPII and TRAPPIII. **B.** The number of deuteron differences for all analysed peptides over the entire deuterium exchange time course for TRAPPIII compared to TRAPPII (n=3). Red points indicate peptides with significant differences (defined as >5%, >0.5 Da, two tailed T-test p value <0.01) in HDX. Only subunits with significant differences between complexes are shown. The full H/D exchange data is summarized in the source data. Increases in the number of deuterons indicates a stabilization in TRAPPII while a decrease in the number of deuterons indicates a stabilization in TRAPPIII. Every point represents the central residue of an individual peptide. **C.** Significant differences in HDX between the TRAPP complexes are mapped on to the core and TRAPPC2L model, with Rab shown to illustrate the binding interface. Increases in exchange are regions that are more stable in TRAPPII, with decreases in exchange representing regions that are more stable in TRAPPIII. Regions with differences are coloured according to the legend. The exact molecular details for how TRAPPC2L associate with the core is unknown, however, it is proposed to bind TRAPPC6, and is positioned on this side of the complex [24].

### The C-terminus of Rab GTPases is not the sole determinant of selectivity for mammalian TRAPP complexes

The conformational differences observed in the TRAPPC4 subunit between TRAPPII and TRAPPIII at a region that interacts with the N-terminus and C-terminus of Rab substrates led us to investigate the potential role of the Rab C-terminus in controlling substrate specificity. We generated two chimeras of both Rab1 and Rab11 where we replaced part of the C-terminal helix and the entire hyper-variable tail (Fig. 5A+B). We conducted GEF assays using TRAPPIII against the four different constructs and found that changing the C-terminus of Rabs did not alter the substrate specificity, with no activity at all against Rab11 with different C-termini, and slightly increased activity with toward Rab1A with the chimeric Rab11A C-termini (Fig. 5C). The hyper-variable tail of Rabs play an essential role in controlling Rab selectivity in the yeast TRAPPII and TRAPPIII complexes, with this mediated by a steric gating mechanism [27]. The chimera data with mammalian TRAPP complexes reveals that there is some determinant of Rab selectivity that is independent of the C-termini and may be driven either by conformational changes in the Rab binding site, or additional interactions between the TRAPPII or TRAPPIII specific subunits and the substrate Rabs.

**Figure 5.**
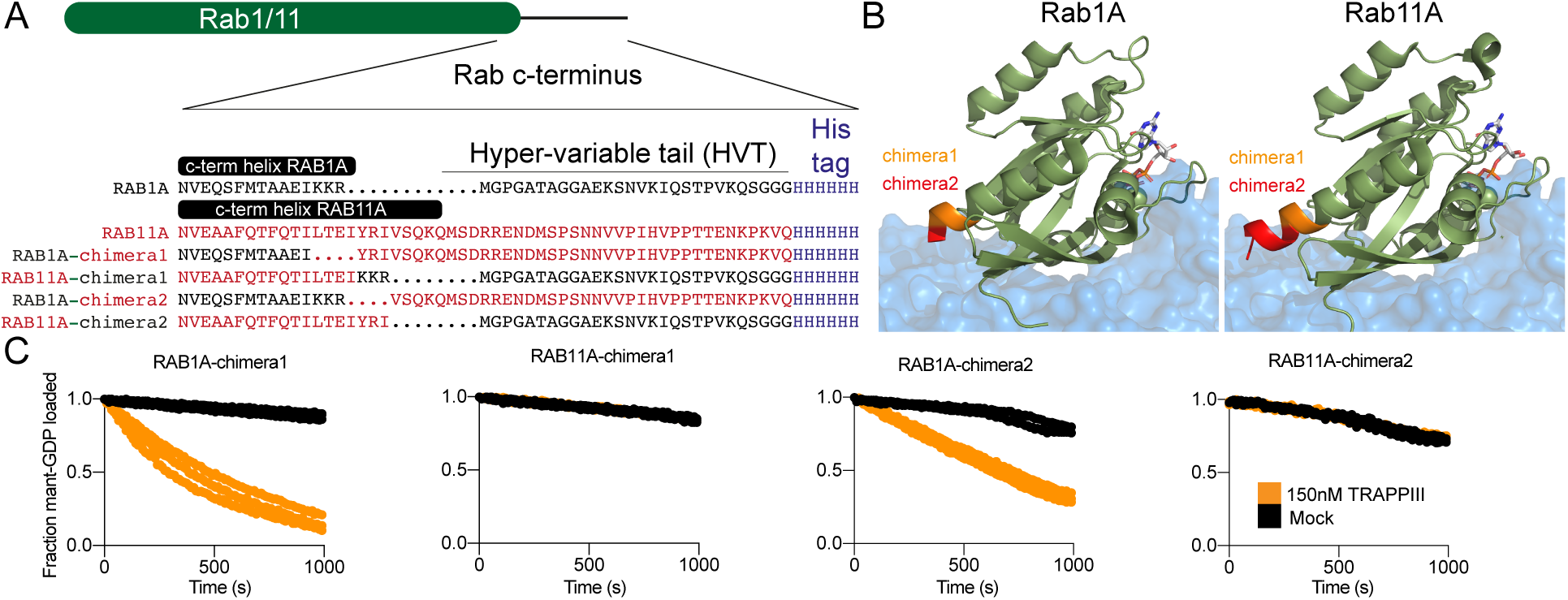
The C-terminus of Rab1A and Rab11 is not the major determinant of Rab selectivity for mammalian TRAPPIII. **A.** Alignment of the C-terminus of Rab1A and Rab11A, covering the C-terminal helix of the Nucleotide binding domain, and the hyper-variable tail. The lipidated cysteine residue is replaced by a his-tag, to allow for Ni-NTA mediated coupling to membranes. Chimeras of Rab1 and Rab11 were generated replacing the C-terminus of Rabs as indicated by the sequence alignment. Rab1A sequence is in black, with the Rab11A sequence in red. **B.** Structures of Rab1A (pdb: 3tkl) [29] and Rab11A (pdb: 6djl) [30] with the approximate position of the TRAPP core as described in Cai et al (pdb: 3cue) [22] The colors indicate the locations where the sequence is swapped for the two chimeras. **C.** In vitro GEF assay of TRAPPIII against Rab1a and Rab11a c-terminal chimeras. 4 *µ*M Rab was loaded with Mant-GDP in the presence of 150 nM TRAPPIII. Control experiments without GEF present were conducted in duplicate for Rab1A and Rab11A chimera1 (n=2), all other concentrations and controls for all other experiments were done in triplicate (n=3).

### Membrane binding enhances GEF activity of TRAPPII and TRAPPIII, and leads to extensive conformational changes

To investigate the role that membranes play in altering TRAPPII and TRAPPIII Rab activity we compared TRAPPII and TRAPPIII mediated GEF activity of Rab1 on extruded liposomes and conducted HDX-MS experiments examining conformational changes that occur upon membrane binding for both TRAPII and TRAPPIII. We tested TRAPPII’s and TRAPPIII’s ability to activate Rab1a on synthetic membranes, with both having enhanced GEF activity. The TRAPPIII complex was more active than TRAPPII with membrane localised Rab1a (43.2 x 10^3^ M^−1^s^−1^ vs 8.8 x 10^3^ M^−1^s^−1^ respectively, Fig. 6A+B). This is consistent with TRAPPIII being the primary activator of Rab1 *in-vivo* and further validates that TRAPP complexes are more efficient GEFs when their substrates are presented on a membrane surface.

**Figure 6.**
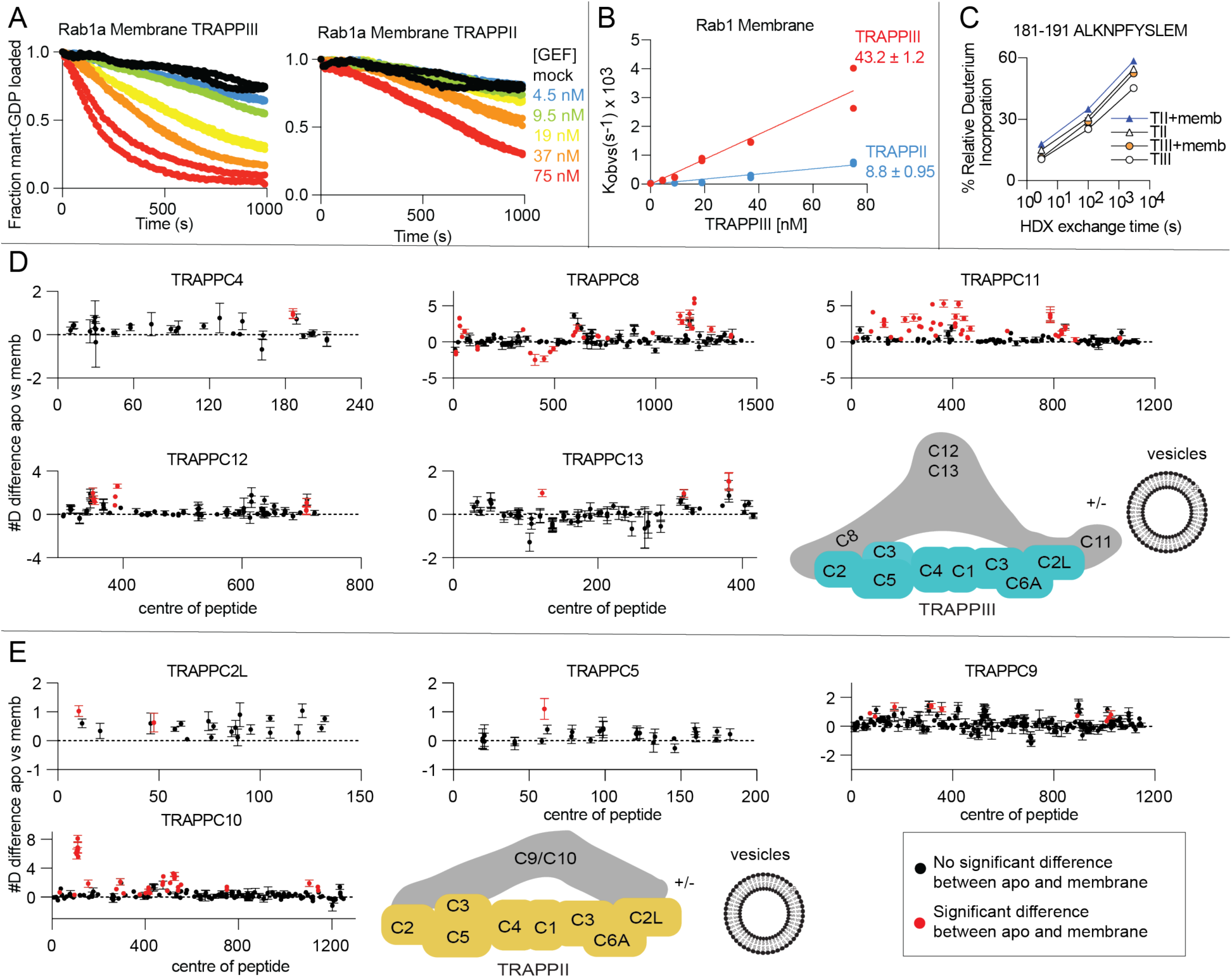
Membrane binding increases GEF activity of both TRAPPII and TRAPPIII, and leads to large scale conformational changes. **A.** In vitro GEF assay of TRAPPIII and TRAPPII at various concentrations (4.5nM – 75nM) against Rab1a (4 *µ*M) in the presence or absence of 150 nm extruded liposomes at 0.2mg/ml (45% PC, 10% PI, 15% PE, 10% PS, 10% DGS NTA, 10% PI(4)P, n=2). **B.** Nucleotide exchange rates of Rab1 plotted as a function of TRAPP concentration (n=2) **C.** Changes in percent deuterium exchange of the TRAPPC4 peptide 181-191 for TRAPPIII (TIII) with and without membrane and TRAPPII (TII) with and without membrane. **D.** The number of deuteron difference for all analysed peptides over the entire deuterium exchange time course for TRAPPIII in the presence and absence of 150 nm extruded liposomes (45% PC, 10% PI, 15% PE, 10% PS, 10% DGS NTA, 10% PI(4)P) (n=3). Only subunits with significant differences are shown (defined as >5%, >0.5 Da, two tailed T-test p value <0.01). Every point represents the central residue of an individual peptide with significant differences indicated in red. **E.** The number of deuteron difference for all analysed peptides over the entire deuterium exchange time course for TRAPPII in the presence and absence of 150 nm extruded liposomes (45% PC, 10% PI, 15% PE, 10% PS, 10% DGS NTA, 10% PI(4)P) (n=3). Only subunits with significant differences are shown (defined as >5%, >0.5Da, two tailed T-test p value <0.01). Every point represents the central residue of an individual peptide with significant differences indicated in red.

Increased GEF activity on membrane surfaces could be mediated by a multitude of mechanisms. Activity could be enhanced by increased local concentration of GEF and substrate on membranes, and/or conformational changes in the TRAPP complex may occur upon membrane binding. We compared differences in HDX-MS upon membrane binding for both TRAPPII and TRAPPIII (Fig. 6D+E). Liposomes were composed of 45% PC, 10% PI, 15% PE, 10% PS, 10% DGS NTA, 10% PI(4)P, mimicking the composition of the trans Golgi network (TGN).

Numerous differences were observed in multiple subunits of both TRAPPII and TRAPPIII in the presence of liposomes (Fig. 6D+E). There was decreased exchange in TRAPPIII in TRAPPC8 (5-21, 115-126, 376-421, 440-457, 471-487, 492-498) and increased exchange was observed in TRAPPC4 (181-191), TRAPPC8 (24-62, 340-350, 588-620, 643-647, 710-720, 759-767, 981-992, 1105-1139, 1159-1198, 1266-1284, 1369-1380), TRAPPC11 (13-25, 74-96, 120-139, 152-161, 193-259, 266-281, 312-338, 349-373, 405-477, 612-621, 773-852, 882-889, 1056-1066), TRAPPC12 (341-358, 382-399, 701-716), and TRAPPC13 (119-127, 311-327, 375-387).

We observed increased exchange in TRAPPII upon membrane binding in TRAPPC2L (8-16, 56-66), TRAPPC5 (51-69), TRAPPC9 (68-80, 87-99, 160-180, 311-322, 352-363, 890-896, 1006-1033) and TRAPPC10 (30-37, 95-124, 149-162, 272-310, 374-381, 397-423, 469-487, 489-562, 740-760, 1093-1112, 1133-1147). The full list of HDX-MS data for all subunits is compiled in the source data. The decreased exchange observed in TRAPPC8 upon membrane binding (5-21, 115-126, 376-421, 440-457, 471-487, 492-498) could be contact sites between TRAPPC8 and membrane or interactions between TRAPPC8 and different subunits that are enhanced in the presence of membranes. The protection in TRAPPC8 (376-421) in the presence of liposomes is consistent with a conserved amphipathic helix present in the yeast TRAPPC8 homolog trs85 (368-409) that was found to be important for membrane binding [23]. This region is highly conserved (Fig. S3) and reveals a key role of TRAPPC8 in driving membrane binding of TRAPPIII.

Increased exchange was observed in the TRAPPIII complex specific subunits (TRAPPC8, TRAPPC11, TRAPPC12, TRAPPC13), and the TRAPPII complex specific subunits (TRAPPC9, TRAPPC10) as well as in the core subunits TRAPPC2L, TRAPPC4, and TRAPPC5 suggesting that large conformational changes occur upon membrane binding. The increased exchange in the core subunits of TRAPPIII (TRAPPC4) include regions that span the Rab binding site (TRAPPC4 181-191) and reveal a possible conformational driven mechanism of activation of TRAPP GEF activity on membranes.

## Discussion

The TRAPPII and TRAPPIII complexes play critical roles in activating Rab GTPases during membrane trafficking processes, with these complexes being conserved in all Eukaryotes. They have well established roles in a diverse set of pathways including secretion, ER-Golgi transport and autophagy [13–16,31]. Many biochemical and structural studies have revealed the architecture of these complexes in both yeast and metazoans showing how these complexes are assembled and how they bind and activate Rab GTPases [14,18,22,32–34]. However, the exact molecular details for TRAPP complex Rab substrate selectivity and membrane activation is still ambiguous. Our biophysical and biochemical analysis of the mammalian TRAPPIII complex has revealed novel insight into TRAPPIII substate specificity, how complex specific subunits alter the Rab binding interface of the core, and how membranes lead to large scale conformational changes in both TRAPPII and TRAPPIII.

The mammalian TRAPPII complex has activity towards Rab1, Rab11a/b, Rab19 and Rab43 [10]. The TRAPPIII complex is a well validated GEF for Rab1a [11,14,15,35], however, the substrate specificity towards other Rab GTPases for mammalian TRAPPIII is unknown. We find that mammalian TRAPPIII has activity towards Rab1a and a novel activity towards Rab43. There was no detectable GEF activity for Rab19 or the other 17 Rab GTPases tested. The observation that TRAPPIII is a GEF for Rab43 but not Rab19 is intriguing since Rab43 and Rab19 are in the Rab1a family and are very evolutionarily similar [28]. TRAPPII was equally active against both Rab43 and Rab19, revealing an additional substrate selectivity between TRAPPII and TRAPPIII. Extensive sequence analysis between Rab1/Rab43 with Rab19/Rab11 does not reveal a simple difference for how TRAPPII and TRAPPIII obtain substrate selectivity. Further biochemical and high resolution structural data will be required to decipher the exact molecular mechanism of Rab specificity.

Multiple conformational differences were observed between TRAPPII and TRAPPIII in the shared subunits. TRAPPIII was more stable at the Rab-binding interface compared with TRAPPII, which suggests initial insight into a possible mechanism for how these complexes can mediate Rab specificity. The C-terminus of TRAPPC4 was more stable in TRAPPIII, with these being very similar regions to those protected upon Rab binding in TRAPPIII and TRAPPII (180-191 in TRAPPC4) [10]. It’s possible that this alteration in the conformation of the putative Rab binding site prevents association with either Rab11 or Rab19 for TRAPPIII. The recent Cryo-EM structure of Drosophila TRAPPIII allowed for a comparison between the two complexes seen in the mammalian complexes [24]. Intriguingly, the interfaces identified between TRAPPC2 with TRAPPC8 and TRAPPC2L with TRAPPC11 identified in the Drosophila TRAPPIII cryo-EM structure were more protected from H/D exchange in TRAPPII compared to TRAPPIII. This suggests that there are shared interfaces between the core and the TRAPPII and TRAPPIII unique subunits. Further details of the differences will require high resolution structural data of the TRAPPII complex.

Rab substrates had increased activity in the presence of synthetic membranes for both TRAPPII and TRAPPIII, consistent with studies of the yeast TRAPPII and TRAPPIII complexes [14]. The TRAPPIII complex activated Rab1a significantly faster in the presence of membranes compared to TRAPPII which is consistent with TRAPPIII being the primary activator of Rab1 *in-vivo*. This increased activity may be driven by the membrane binding site identified in HDX-MS studies in TRAPPC8. We found multiple peptides with significant protections spanning TRAPPC8 including the region 376-421. This region corresponds to an amphipathic helix (368-409) discovered in yeast Trs85, where mutants of this region decreased membrane recruitment and Rab activation [23]. This highlights a conserved role throughout evolution of the TRAPPC8 subunit in mediating membrane association and activation of Rab1. Multiple TRAPPII and TRAPPIII specific subunits were destabilised upon membrane binding. These increases in exchange are likely driven by conformational rearrangements of the subunit-subunit interacting regions. Further detailed structural studies will help interpret how these changes may mediate membrane binding or alter GEF activity.

The emerging set of clinical mutations in either TRAPP core or complex specific subunits demonstrates the important roles that both TRAPPII and TRAPPIII play in membrane trafficking [5,6,9,36]. Interestingly, our HDX-MS experiments revealed conformational differences between TRAPPII and TRAPPIII located at or near these sites of mutations. These differences occurred in TRAPPC2 and TRAPPC2L. The mutations D47Y, H80R, and F83S in TRAPPC2 have been associated with the skeletal disorder spondyloepiphyseal dysplasia tarda (SEDT; also abbreviated SEDL) [37,38]. TRAPPII was protected compared to TRAPPIII in peptides spanning 47-53 and 59-70 of TRAPPC2. The D47Y mutation has been shown to disrupt both the TRAPPIII and TRAPPII complexes through disrupting the interactions between TRAPPC2 with TRAPPC9 or TRAPPC8 [37] with the equivalent D46Y mutation in yeast only disrupting the TRAPPIII complex [38]. The mutation D37Y in TRAPPC2L has been associated with neurodevelopmental delay [39]. The equivalent yeast mutant (D45Y) disrupts the interaction between TRAPPC2L and the TRAPPII specific subunit TRAPPC10 [39]. We found large stabilisations in TRAPPII compared to TRAPPIII in a peptide spanning 37-58 which would be consistent with this mutation disrupting the TRAPPC2L-TRAPPC10 interaction. These differences between TRAPPII and TRAPPIII at these important residues indicate that the clinical mutants could possibly target TRAPPII and TRAPPIII differently depending on unique interactions with complex specific subunits.

HDX-MS experiments examining differences upon membrane binding revealed secondary structure differences at or near regions mutated in human disease, specifically in TRAPPC9 and TRAPPC11. The mutation L178P in TRAPPC9 has been implicated with severe intellectual disability [40]. HDX-MS experiments comparing apo-TRAPPII to membrane bound TRAPPII showed an exposure in a peptide spanning 160-180. Furthermore, the mutations Q284P and Q777P of TRAPPC11 have been implicated in muscular disorders by elevated creatine kinase levels and cerebral atrophy [41,42] and we found that TRAPPC11 was exposed in the peptides 266-281 and 773-801. The peptide spanning 266-281 lies extremely close to the Q284 residue and the peptide spanning 773-801 completely contains the Q777 residue. These secondary structure differences could indicate that these clinical mutants alter the membrane activation of both TRAPPII and TRAPPIII complexes. More research is needed to investigate the exact molecular mechanisms of these clinical mutants and how they alter TRAPP’s function.

The TRAPP complexes are some of the most important regulators of membrane trafficking and activators of Rabs at the Golgi. Our data highlights the critical dynamic differences between the two complexes at the Rab binding site which likely has a role in regulating Rab specificity. Continued biochemical and structural studies will be required to decipher the exact mechanism of this specificity with this work helping provide a strong start point for these future studies.

## Acknowledgements

This work in the Burke lab was supported by Natural Science and Engineering Research Council (NSERC Discovery Grant 2020-04241), and a Michael Smith Foundation for Health Research Scholar award (17868). C.K.Y. is supported by CIHR (FDN-143228) and the Natural Sciences and Engineering Research Council of Canada (RGPIN-2018-03951). This research project was supported in part by the UBC High Resolution Macromolecular Cryo-Electron Microscopy Facility (HRMEM).

## Contributions

JEB, NH and MLJ designed all biophysical/biochemical experiments. NH cloned the TRAPPIII complex. NH, MP and MLJ carried out protein expression/purification. NH and MLJ carried out all biochemical studies. NH, MLJ, KDF and MP carried out HDX-MS experiments. UD SEN and CKY carried out electron microscopy studies. NH, MLJ, and JEB wrote the manuscript with input from all authors.

## Materials and Methods

### Plasmids and antibodies

The full length TRAPPII, and Rab genes were used as described previously [10]. TRAPPC8, TRAPPC11, and TRAPPC12 genes, were purchased from DNASU (C8-HsCD00347731 & HsCD00399392, C11-HsCD00082480, C12-HsCD324976) and TRAPPC13 was ordered from Thermofisher Geneart. Genes were subcloned into pLIB vectors, and in the case of TRAPPC12 a TEV cleavable c-term 2x strep tag was added while a TEV cleavable c-term 6x his tag was added to the c-term of TRAPPC11. Genes were subsequently amplified following the biGBac protocol to generate 2 plasmids that together contain all the TRAPPIII genes [43]. A table summarizing the plasmids used is outlined in Table S1.

### Protein expression

All TRAPPII complexes were similarly expressed as previously described [10]. In short, to express TRAPPII/TRAPPIII complexes, an optimized ratio of baculovirus was used to co-infect *Spodoptera frugiperda* (Sf9) cells between 1-2×10^6^ cells/mL. Co-infections were harvested at ∼66-hours and washed with ice-cold PBS before snap-freezing in liquid nitrogen. Rab constructs were all expressed in BL21 C41 *E.coli,* induced with 0.5mM IPTG and grown at 37°C for 4hrs. Pellets were washed with ice-cold phosphate-buffered saline (PBS), flash frozen in liquid nitrogen, and stored at −80°C until use.

### Protein purification

TRAPPII complexe and Rabs were purified as described previously [10]. TRAPP cell pellets were lysed by sonication for 1.5 minutes in lysis buffer (20mM Tris pH 8.0, 100mM NaCl, 5% (v/v) glycerol, 2mM ß–mercaptoethanol (BME), and protease inhibitors (Millipore Protease Inhibitor Cocktail Set III, Animal-Free)). Triton X-100 was added to 0.1% v/v, and the solution was centrifuged for 45 minutes at 20,000 x g at 1°C. The supernatant was then loaded onto a 5 mL HisTrap™ FF column (GE Healthcare) that had been equilibrated in NiNTA A buffer (20 mM Tris pH8.0, 100 mM NaCl, 10 mM imidazole pH 8.0, 5% (v/v) glycerol, 2 mM bME). The column was washed with 20 mL of NiNTA buffer, 20 mL of 6% NiNTA B buffer (20 mM Tris pH 8.0, 100 mM NaCl, 200 mM imidazole pH 8.0, 5% (v/v) glycerol, 2 mM bME) before being eluted with 100% NiNTA B. The eluate was subsequently loaded on a 5ml Strep™ column and washed with 10ml SEC buffer (20mM HEPES pH 7.5, 100mM NaCl, 0.5mM TCEP). The Strep-tag was cleaved by adding SEC buffer containing 10mM BME and TEV protease to the column and incubating overnight at 4°C. Protein was pooled and concentrated using Amicon 50K concentrator and size exclusion chromatography (SEC) was performed using a Superose 6 increase 10/300 column equilibrated in SEC Buffer. Yield can be improved by taking pooled TRAPP complex before SEC and manually running protein through a 1 mL HisTrap™ FF column (GE Healthcare) that had been equilibrated in NiNTA A buffer followed by buffer exchange into SEC buffer using a 5 mL HiTrap® Desalting Column (GE Healthcare). Fractions containing protein of interest were pooled, concentrated, flash frozen in liquid nitrogen and stored at −80°C.

For Rab purification, cell pellets were lysed by sonication for 5 minutes in lysis buffer (20mM Tris pH 8.0, 100mM NaCl, 5% (v/v) glycerol, 2mM ß–mercaptoethanol (BME), and protease inhibitors (Millipore Protease Inhibitor Cocktail Set III, Animal-Free)). Triton X-100 was added to 0.1% v/v, and the solution was centrifuged for 45 minutes at 20,000 x g at 1°C. Supernatant was loaded onto a 5ml GSTrap 4B column (GE Healthcare) in a superloop for 1.5 hours and the column was washed in Buffer A (20mM Tris pH 8.0, 100mM NaCl, 5% (v/v) glycerol, 2mM BME) to remove non-specifically bound proteins. The GST-tag was cleaved by adding Buffer A containing 10mM BME and TEV protease to the column and incubating overnight at 4°C. Cleaved protein was eluted with Buffer A. Protein was further purified by separating on a 5ml HiTrap Q column with a gradient of Buffer A and Buffer B (20mM Tris pH 8.0, 1M NaCl, 5% (v/v) glycerol, 2mM BME). Protein was pooled and concentrated using an Amicon 30K concentrator, and were flash frozen in liquid nitrogen and stored at −80°C.

### Lipid vesicle preparation

Nickelated lipid vesicles were made with 10% phosphatidylserine (bovine brain PS, Sigma), 10% L-α-phosphatidylinositol-4-phosphate (PI4P, Avanti) 30% phosphatidylcholine (egg yolk PC, Sigma), 10% L-α-Phosphatidylinositol (Liver PI, Avanti), 15% L-α-Phosphatidylethanolamine (egg yolk PE, Sigma), and 10% DGS-NTA(Ni) (18:1 DGSNTA(Ni), Avanti). Vesicles were prepared by combining liquid chloroform stocks together at appropriate concentrations and evaporating away the chloroform with nitrogen gas. The resulting lipid film layer was desiccated for 20 min before being resuspended in lipid buffer (20mM HEPES (pH 7.5) and 100mM KCl) to a concentration of 1mg/ml. The lipid solution was vortexed for 5 min, bath sonicated for 10 min, and flash frozen in liquid nitrogen. Vesicles were then subjected to three freeze thaw cycles using a warm water bath. Vesicles were extruded 11 times through a 150nm NanoSizer Liposome Extruder (T&T Scientific) and stored at −80°C.

### In-vitro GEF assay

C-terminally His-tagged Rabs were purified, and nucleotide loaded as described previously [10]. Reactions were conducted in 10*µ*l volumes with a final concentration of 4 *µ*M Mant-GDP loaded Rab, 100 *µ*M GTPγS, the appropriate amount of TRAPP (4.5 – 300 *µ*M) and synthetic vesicles (0.2mg/ml). Rab and membrane were aliquoted into a 384-well, black, low-volume plate (Corning 3676). To start the reaction, TRAPP and GTPγS were added simultaneously to the wells and a SpectraMax® M5 Multi-Mode Microplate Reader was used to measure the fluorescent signal for 1hr (Excitation λ = 366nm; Emission λ = 443nm). Data was analyzed using GraphPad Prism 7 Software, and k_cat_/K_m_ analysis was carried out according to the protocol of [44]. GEF curves were fit to a non-linear dissociate one phase exponential decay using the formula I(t)=(I_0_-I_∞_)*exp(-k_obs_*) + I_∞_ (GraphPad Software), where I(t) is the emission intensity as a function of time, and I_0_ and I_∞_ are the emission intensities at t=o and t=∞. The catalytic efficiency k_cat_/K_m_ was obtained by a slope of a linear least squares fit to k_obs_=k_cat_/K_m_*[GEF]+ k_intr_, where k_intr_ is the rate constant in the absence of GEF.

### Hydrogen deuterium exchange (HDX)

#### TRAPPII vs TRAPPIII

HDX reactions comparing both complexes were conducted in 50*µ*l reaction volumes with a final concentration of 160nM for TRAPPII or TRAPPIII. Exchange was carried out in triplicate for five time points (3s at 4°C and 3s, 30s, 300s and 3000s at 20°C). Hydrogen deuterium exchange was initiated by the addition of 40*µ*l of D_2_O buffer solution (10mM HEPES pH 7.5, 50mM NaCl, 97% D_2_O) to give a final concentration of 78% D_2_O. Exchange was terminated by the addition of acidic quench buffer at a final concentration 0.6M guanidine-HCl and 0.9% formic acid. Samples were immediately frozen in liquid nitrogen at −80°C.

### Hydrogen deuterium exchange (HDX) TRAPPII and TRAPPIII apo vs Membrane

HDX reactions comparing both complexes bound to membranes were conducted in 50*µ*l reaction volumes with a final concentration of 120nM for TRAPPII or TRAPPIII and 0.1mg/ml membranes. Exchange was carried out in triplicate for three time points (3s, 100s and 3000s at 20°C). Prior to the addition of D_2_O, TRAPP was incubated at 20°C with vesicles for one minute to facilitate TRAPP-membrane interactions. Hydrogen deuterium exchange was initiated by the addition of 32.1*µ*l of D_2_O buffer solution (10mM HEPES pH 7.5, 50mM NaCl, 97% D_2_O) to the protein/membrane solutions, to give a final concentration of 62% D_2_O. Exchange was terminated by the addition of acidic quench buffer at a final concentration 0.6M guanidine-HCl and 0.9% formic acid. Samples were immediately frozen in liquid nitrogen at −80°C.

### Hydrogen deuterium exchange (HDX) TRAPPIII apo vs Rab bound

HDX reactions comparing TRAPPIII and TRAPPIII-Rab1 complex were conducted in 50*µ*l reaction volumes with a final concentration 100nM TRAPPIII per sample and 560 nM Rab1. Exchange was carried out in triplicate for four time points (3s, 30s, 300s and 3000s at 20°C). Prior to the addition of D_2_O, proteins were incubated on ice in the presence of 20mM EDTA for 30 minutes to facilitate release of nucleotide. Hydrogen deuterium exchange was initiated by the addition of 43.1*µ*l of D_2_O buffer solution (10mM HEPES pH 7.5, 50mM NaCl, 97% D_2_O) to give a final concentration of 82% D_2_O. Exchange was terminated by the addition of acidic quench buffer at a final concentration 0.6M guanidine-HCl and 0.9% formic acid. Samples were immediately frozen in liquid nitrogen at −80°C.

### HDX-MS data Analysis

Protein samples were rapidly thawed and injected onto an integrated fluidics system containing a HDx-3 PAL liquid handling robot and climate-controlled (2°C) chromatography system (LEAP Technologies), a Dionex Ultimate 3000 UHPLC system, as well as an Impact HD QTOF Mass spectrometer (Bruker) [45]. The protein was run over one (at 10°C) immobilized pepsin column (Trajan; ProDx protease column, 2.1 mm x 30 mm PDX.PP01-F32) at 200 *µ*L/min for 3 minutes. The resulting peptides were collected and desalted on a C18 trap column (Acquity UPLC BEH C18 1.7mm column (2.1 x 5 mm); Waters 186003975). The trap was subsequently eluted in line with an ACQUITY 1.7 μm particle, 100 × 1 mm2 C18 UPLC column (Waters), using a gradient of 3-35% B (Buffer A 0.1% formic acid; Buffer B 100% acetonitrile) over 11 minutes immediately followed by a gradient of 35-80% over 5 minutes. Mass spectrometry experiments acquired over a mass range from 150 to 2200 m/z using an electrospray ionization source operated at a temperature of 200C and a spray voltage of 4.5 kV. The resulting MS/MS datasets were analyzed using PEAKS7 (PEAKS), and a false discovery rate was set at 1% using a database of purified proteins and known contaminants [46]. HDExaminer Software (Sierra Analytics) was used to automatically calculate the level of deuterium incorporation into each peptide. All peptides were manually inspected for correct charge state and presence of overlapping peptides. Deuteration levels were calculated using the centroid of the experimental isotope clusters. Differences in exchange were in a peptide were considered significant if they met all three of the following criteria: >5% change in exchange, >0.5 Da difference in exchange, and a p value <0.01 using a two tailed student t-test. Samples were only compared within a single experiment and were never compared to experiments completed at a different time with a different final D_2_O level.

The data analysis statistics for all HDX-MS experiments are in supplemental tables 2-4 according to the guidelines of [47]. The mass spectrometry proteomics data have been deposited to the ProteomeXchange Consortium via the PRIDE partner repository [48] with the dataset identifier PXD025928.

### Negative stain single-particle electron microscopy (EM) and image analysis

Purified TRAPPIII complex was adsorbed to glow discharged carbon coated grids and stained with uranyl formate. The stained specimens were examined using a Talos L120C transmission electron microscope (ThermoFisher Scientific) operated at an accelerating voltage of 120 kV and equipped with a Ceta charged-coupled-device (CCD) camera. 100 micrographs were acquired at a nominal magnification of 45,000x at a defocus of ∼1.2μm and binned twice to obtain a final pixel size of 4.53 Å/pixel. Contrast transfer function (CTF) estimation for each micrograph was carried out using CTFFind4 [49]. 200 particles were manually picked then aligned to generate 2D class averages for template-based autopicking in Relion 3.0 [50]. 38,062 particles were autopicked and extracted with a box size of 148 pixels. Particles were then subjected to 2D classification and 1034 particles which did not classify well were discarded. The remaining particles were transferred to cryoSPARC v2.14 [51] for *ab initio* reconstruction using 2 classes which were refined by heterogenous refinement. A final particle stack of 32,429 particles was used to carry out homogenous refinement of the better model, yielding a final map at 14.2Å resolution based on the gold-standard 0.143 Fourier Shell Correlation criterion (Fig. S1). The EM data have been deposited to the EMDB with the accession code: EMD-23997.

## Supplemental Information for

**Supplemental Figure 1.**
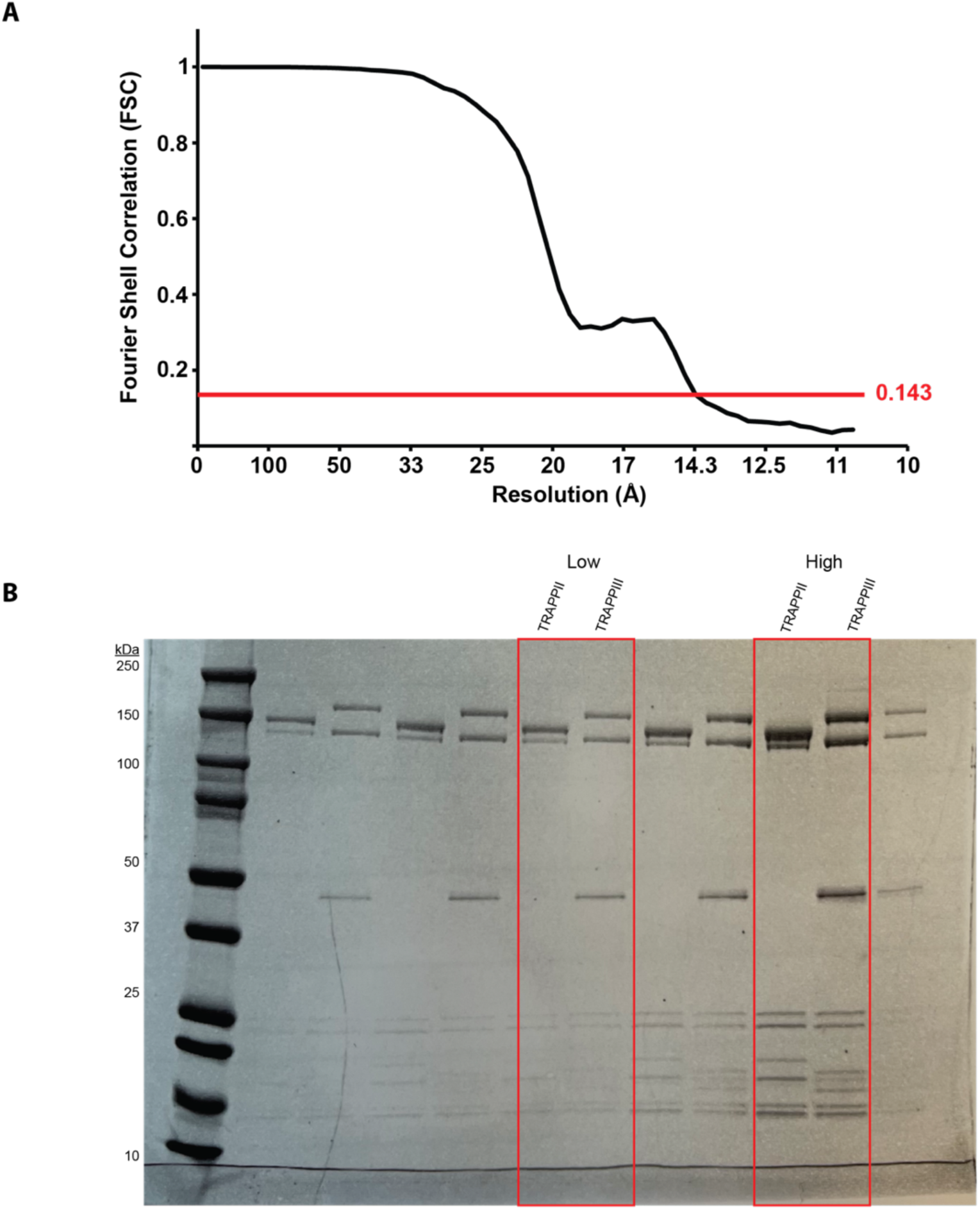
Gold-standard Fourier shell correlation and SDS-PAGE gel of purified TRAPP complexes. (A) Gold-standard Fourier shell correlation curve showing the resolution of the TRAPPIII model in figure 1. (B) Uncropped SDS-PAGE gel of TRAPPII and TRAPPIII used in figure 1. High and low refer to protein amount loaded (high=3.6ug and low= 1.4ug). 4-20% NuPAGE gradient gel run at 225V for 35 min and stained with Coomassie Brilliant Blue dye.

**Supplemental Figure 2.**
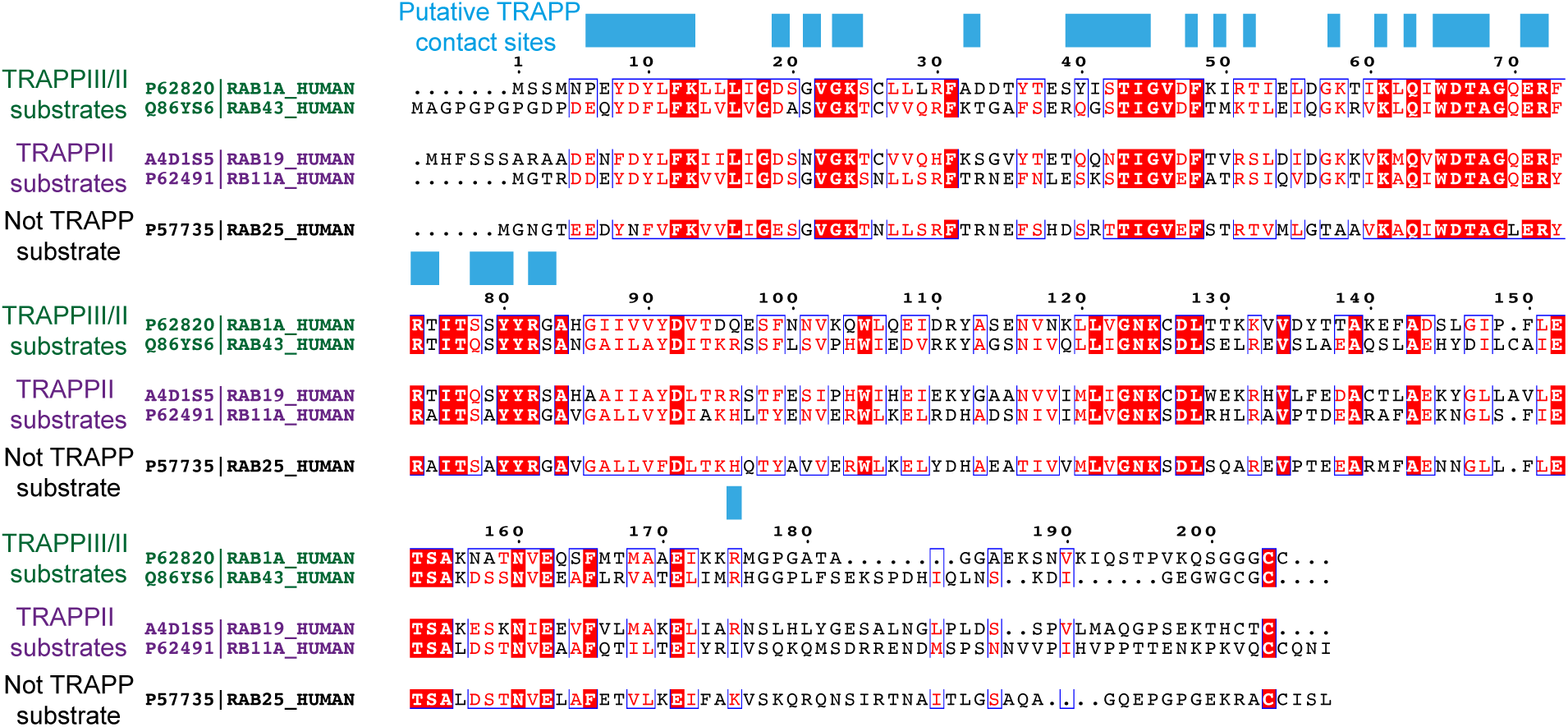
Alignment of substrate Rabs for mammalian TRAPP complexes. Alignment of Rab1, Rab43, Rab19, Rab11a, and Rab25. Putative contact sites with the TRAPP core are based on the pdb: 3CUE. Alignment was generated using Espript [1].

**Supplemental Figure 3.**
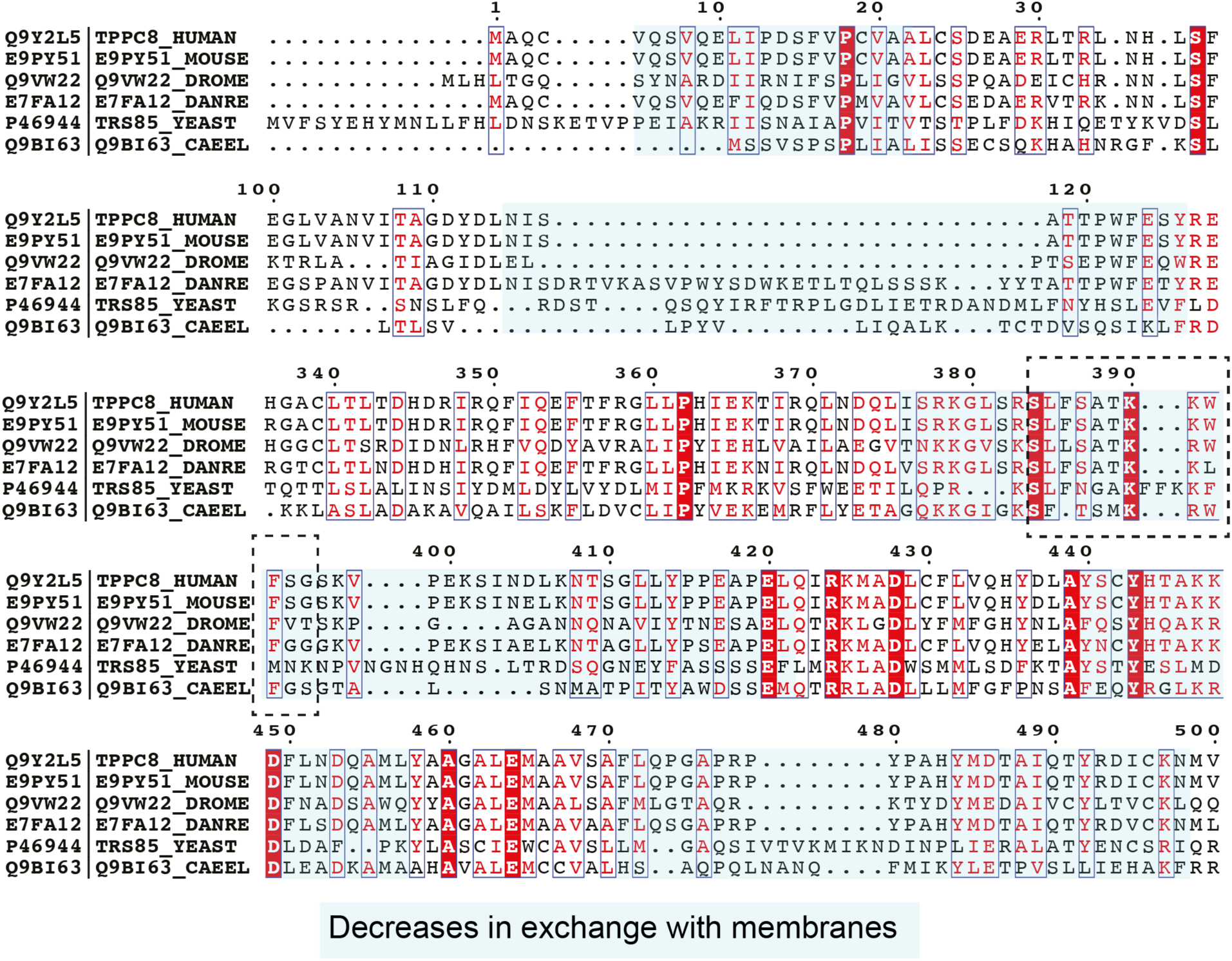
Alignment of TRAPPC8. Decreases in exchange upon membrane binding are highlighted in light blue. The putative amphipathic helix from Trs85 is indicated in the boxed dotted line. Alignment was generated using Espript [1].

**Supplemental table 1.** All plasmids used in this study.

| Name | Plasmid | Protein(s) | Sequence(s) | Modifications/Tags | Source |
| --- | --- | --- | --- | --- | --- |
| MJ153<br>[TRAPPIIa] | pBIG1a | TRAPPC1;<br>TRAPPC2L;<br>TRAPPC5;<br>TRAPPC6a;<br>TRAPPC10; | 1-145;<br>1-140;<br>1-188;<br>1-173;<br>1-1259 | TRAPPC10 – C-term His Tag (TEV) | [2] |
| MJ85<br>[TRAPPIIb] | pBIG1a | TRAPPC3;<br>TRAPPC2;<br>TRAPPC4;<br>TRAPPC9 | 1-180;<br>1-140;<br>1-235;<br>1-1148 | TRAPPC3 – C-term Strep Tag (TEV) | [2] |
| NH40<br>[TRAPPIIIa] | pBIG1a | TRAPPC8;<br>TRAPPC11;<br>TRAPPC12;<br>TRAPPC13 | 1-1435;<br>1-1147;<br>300-735;<br>1-417 | TRAPPC12 - C-term Strep Tag (TEV);<br>TRAPPC11 - C-term His Tag (TEV) | This paper |
| NH42<br>[TRAPPIIIb] | pBIG2ab | TRAPPC1;<br>TRAPPC2;<br>TRAPPC2L;<br>TRAPPC3;<br>TRAPPC4;<br>TRAPPC5;<br>TRAPPC6a | 1-145;<br>1-140;<br>1-140;<br>1-180;<br>1-235;<br>1-188;<br>1-173; |  | This paper |
| MJ117 | pOPTGcH | Rab1a | 1-203 | N-term GST Tag(TEV); C-term His Tag | [2] |
| MJ37 | pOPTGcH | Rab2a | 1-210 | N-term GST Tag(TEV); C-term His Tag | [2] |
| EM11 | pOPTGcH | Rab3a | 1-217 | N-term GST Tag(TEV); C-term His Tag | [2] |
| MJ36 | pOPTGcH | Rab4b | 1-210 | N-term GST Tag(TEV); C-term His Tag | [2] |
| DH1 | pOPTGcH | Rab5a | 1-212 | N-term GST Tag(TEV); Q79L mutation | [2] |
| EM13 | pOPTGcH | Rab6a | 1-205 | N-term GST Tag(TEV); C-term His Tag | [2] |
| NH20 | pOPTGcH | Rab7a | 1-204 | N-term GST Tag(TEV); C-term His Tag | [2] |
| MJ40 | pOPTGcH | Rab8a | 1-203 | N-term GST Tag(TEV); C-term His Tag | [2] |
| EO7 | pOPTGcH | Rab11a | 1-211 | N-term GST Tag(TEV); C-term His Tag | [2] |
| MJ28 | pOPTGcH | Rab11b | 1-213 | N-term GST Tag(TEV); C-term His Tag | [2] |
| MJ39 | pOPTGcH | Rab12 | 1-242 | N-term GST Tag(TEV); C-term His Tag | [2] |
| MJ38 | pOPTGcH | Rab14 | 1-212 | N-term GST Tag(TEV); C-term His Tag | [2] |
| EM15 | pOPTGcH | Rab18 | 1-198 | N-term GST Tag(TEV); C-term His Tag | [2] |
| MJ167 | pOPTGcH | Rab19 | 1-214 | N-term GST Tag(TEV); C-term His Tag | [2] |
| MJ27 | pOPTGcH | Rab25 | 1-208 | N-term GST Tag(TEV); C-term His Tag | [2] |
| NH19 | pOPTGcH | Rab29 | 1-201 | N-term GST Tag(TEV); C-term His Tag | [2] |
| NH21 | pOPTGcH | Rab32 | 1-223 | N-term GST Tag(TEV); C-term His Tag | [2] |
| EM17 | pOPTGcH | Rab33a | 1-234 | N-term GST Tag(TEV); C-term His Tag | [2] |
| Em19 | pOPTGcH | Rab35 | 1-199 | N-term GST Tag(TEV); C-term His Tag | [2] |
| EM23 | pOPTGcH | Rab39a | 1-214 | N-term GST Tag(TEV); C-term His Tag | [2] |
| EM25 | pOPTGcH | Rab43 | 1-209 | N-term GST Tag(TEV); C-term His Tag | [2] |
| MJ210 | pOPTGcH | Rab11A chimera 1 | 1-209 | N-term GST Tag(TEV); C-term His Tag;<br>Rab11 tail swapped with Rab1 tail | This paper |
| MJ211 | pOPTGcH | Rab1A chimera 1 | 1-217 | N-term GST Tag(TEV); C-term His Tag;<br>Rab1a tail swapped with Rab11a tail | This paper |
| MJ201 | pOPTGcH | Rab11A chimera 2 | 1-209 | N-term GST Tag(TEV); C-term His Tag;<br>Rab11a tail swapped with Rab1 tail | This paper |
| MJ202 | pOPTGcH | Rab1A chimera 2 | 1-217 | N-term GST Tag(TEV); C-term His Tag;<br>Rab1a tail swapped with Rab11 tail | This paper |

**Supplemental Table 2.** HDX Statistics for Figure 3 - TRAPPIII Rab.

| Data set | TRAPP C1 | TRAPP C2 | TRAPP C2L | TRAPP C3 | TRAPP C4 | TRAPP C5 |
| --- | --- | --- | --- | --- | --- | --- |
| HDX reaction details | %D.O=82%<br>pH<sub>int</sub>=7.5<br>Temp=20°C | %D.O=82%<br>pH<sub>int</sub>=7.5<br>Temp=20°C | %D.O=82%<br>pH<sub>int</sub>=7.5<br>Temp=20°C | %D.O=82%<br>pH<sub>int</sub>=7.5<br>Temp=20°C | %D.O=82%<br>pH<sub>int</sub>=7.5<br>Temp=20°C | %D.O=82%<br>pH<sub>int</sub>=7.5<br>Temp=20°C |
| HDX time course (s) | 3, 30, 300, 3000 | 3, 30, 300, 3000 | 3, 30, 300, 3000 | 3, 30, 300, 3000 | 3, 30, 300, 3000 | 3, 30, 300, 3000 |
| HDX controls | N/A | N/A | N/A | N/A | N/A | N/A |
| Back-exchange | Corrected based on %D.O | Corrected based on %D.O | Corrected based on %D.O | Corrected based on %D.O | Corrected based on %D.O | Corrected based on %D.O |
| Number of peptides | 18 | 25 | 22 | 24 | 29 | 23 |
| Sequence coverage | 97.2 | 87.1 | 90.7 | 58.3 | 96.3 | 87.2 |
| Average peptide /redundancy | Length=11.9<br>Redundancy=1.5 | Length=13.3<br>Redundancy=2.4 | Length=11.9<br>Redundancy=1.9 | Length=10.6<br>Redundancy=1.4 | Length=12.2<br>Redundancy=1.6 | Length=13<br>Redundancy=1.6 |
| Replicates | 3 | 3 | 3 | 3 | 3 | 3 |
| Repeatability | Average<br>StDev=0.9% | Average<br>StDev=0.7% | Average<br>StDev=1.4% | Average<br>StDev=1.1% | Average<br>StDev=1% | Average<br>StDev=1.1% |
| Significant differences in HDX | >5% and >0.5 Da and unpaired t-test ≤0.01 | >5% and >0.5 Da and unpaired t-test ≤0.01 | >5% and >0.5 Da and unpaired t-test ≤0.01 | >5% and >0.5 Da and unpaired t-test ≤0.01 | >5% and >0.5 Da and unpaired t-test ≤0.01 | >5% and >0.5 Da and unpaired t-test ≤0.01 |
| Data set | TRAPP C6 | TRAPP C8 | TRAPP C11 | TRAPP C12 | TRAPP C13 | Rab1a |
| HDX reaction details | %D.O=82%<br>pH<sub>int</sub>=7.5<br>Temp=20°C | %D.O=82%<br>pH<sub>int</sub>=7.5<br>Temp=20°C | %D.O=82%<br>pH<sub>int</sub>=7.5<br>Temp=20°C | %D.O=82%<br>pH<sub>int</sub>=7.5<br>Temp=20°C | %D.O=82%<br>pH<sub>int</sub>=7.5<br>Temp=20°C | %D.O=82%<br>pH<sub>int</sub>=7.5<br>Temp=20°C |
| HDX time course (seconds) | 3, 30, 300, 3000 | 3, 30, 300, 3000 | 3, 30, 300, 3000 | 3, 30, 300, 3000 | 3, 30, 300, 3000 | 3, 30, 300, 3000 |
| HDX controls | N/A | N/A | N/A | N/A | N/A | N/A |
| Back-exchange | Corrected based on %D.O | Corrected based on %D.O | Corrected based on %D.O | Corrected based on %D.O | Corrected based on %D.O | Corrected based on %D.O |
| Number of peptides | 21 | 157 | 126 | 50 | 84 | 41 |
| Sequence coverage | 73.4 | 89 | 77.2 | 82.7 | 95.7 | 87.3 |
| Average peptide /redundancy | Length=9.2<br>Redundancy=1.1 | Length=15.1<br>Redundancy=1.7 | Length=12.4<br>Redundancy=1.4 | Length=13.7<br>Redundancy=1.6 | Length=11.2<br>Redundancy=2.2 | Length=14<br>Redundancy=2.8 |
| Replicates | 3 | 3 | 3 | 3 | 3 | 3 |
| Repeatability | Average<br>StDev=1.4% | Average<br>StDev=0.9% | Average<br>StDev=0.9% | Average<br>StDev=0.9% | Average<br>StDev=0.9% | Average<br>StDev=0.9% |
| Significant differences in HDX | >5% and >0.5 Da and unpaired t-test ≤0.01 | >5% and >0.5 Da and unpaired t-test ≤0.01 | >5% and >0.5 Da and unpaired t-test ≤0.01 | >5% and >0.5 Da and unpaired t-test ≤0.01 | >5% and >0.5 Da and unpaired t-test ≤0.01 | >5% and >0.5 Da and unpaired t-test ≤0.01 |

**Supplemental Table 3.** HDX Statistics for Figure 4 - TRAPPII vs TRAPPIII.

| Data set | TRAPP C1 | TRAPP C2 | TRAPP C2L | TRAPP C3 | TRAPP C4 | TRAPP C5 | TRAPP C6 |
| --- | --- | --- | --- | --- | --- | --- | --- |
| HDX reaction details | %D.O.=78%<br>pH<sub>in</sub>=7.5<br>Temp=4-20°C | %D.O.=78%<br>pH<sub>in</sub>=7.5<br>Temp=4-20°C | %D.O.=78%<br>pH<sub>in</sub>=7.5<br>Temp=4-20°C | %D.O.=78%<br>pH<sub>in</sub>=7.5<br>Temp=4-20°C | %D.O.=78%<br>pH<sub>in</sub>=7.5<br>Temp=4-20°C | %D.O.=78%<br>pH<sub>in</sub>=7.5<br>Temp=4-20°C | %D.O.=78%<br>pH<sub>in</sub>=7.5<br>Temp=4-20°C |
| HDX time course (s) | 3 at 4 °C<br>3, 30, 300,<br>3000 at 20°C | 3 at 4 °C<br>3, 30, 300,<br>3000 at 20°C | 3 at 4 °C<br>3, 30, 300,<br>3000 at 20°C | 3 at 4 °C<br>3, 30, 300,<br>3000 at 20°C | 3 at 4 °C<br>3, 30, 300,<br>3000 at 20°C | 3 at 4 °C<br>3, 30, 300,<br>3000 at 20°C | 3 at 4 °C<br>3, 30, 300,<br>3000 at 20°C |
| HDX controls | N/A | N/A | N/A | N/A | N/A | N/A | N/A |
| Back-exchange | Corrected based on % D.O | Corrected based on %D.O | Corrected based on %D.O | Corrected based on %D.O | Corrected based on %D.O | Corrected based on %D.O | Corrected based on %D.O |
| Number of peptides | 22 | 29 | 27 | 27 | 35 | 39 | 25 |
| Sequence coverage | 96.6 | 96.4 | 94.3 | 69.4 | 96.3 | 88.3 | 85.5 |
| Average peptide /redundancy | Length=14.5<br>Redundancy=2.2 | Length= 13.4<br>Redundancy=2.8 | Length=13.6<br>Redundancy= 2.6 | Length=12<br>Redundancy= 2.1 | Length=12.5<br>Redundancy= 2.0 | Length=13.5<br>Redundancy= 2.8 | Length=12.1<br>Redundancy= 1.8 |
| Replicates | 3 | 3 | 3 | 3 | 3 | 3 | 3 |
| Repeatability | Average StDev=0.6% | Average StDev=0.6% | Average StDev=0.9% | Average StDev=0.8% | Average StDev=0.7% | Average StDev=0.8% | Average StDev=0.7% |
| Significant differences in HDX | >5% and >0.5 Da and unpaired t-test ≤0.01 | >5% and >0.5 Da and unpaired t-test ≤0.01 | >5% and >0.5 Da and unpaired t-test ≤0.01 | >5% and >0.5 Da and unpaired t-test ≤0.01 | >5% and >0.5 Da and unpaired t-test ≤0.01 | >5% and >0.5 Da and unpaired t-test ≤0.01 | >5% and >0.5 Da and unpaired t-test ≤0.01 |

**Supplemental Table 4.** HDX Statistics for Figure 6 – Membrane.

| Data set | TRAPP C1 | TRAPP C2 | TRAPP C2L | TRAPP C3 | TRAPP C4 | TRAPP C5 | TRAPP C6 |
| --- | --- | --- | --- | --- | --- | --- | --- |
| HDX reaction details | %D.O=62%<br>pH<sub>int</sub>=7.5<br>Temp=20°C | %D.O=62%<br>pH<sub>int</sub>=7.5<br>Temp=20°C | %D.O=62%<br>pH<sub>int</sub>=7.5<br>Temp=20°C | %D.O=62%<br>pH<sub>int</sub>=7.5<br>Temp=20°C | %D.O=62%<br>pH<sub>int</sub>=7.5<br>Temp=20°C | %D.O=62%<br>pH<sub>int</sub>=7.5<br>Temp=20°C | %D.O=62%<br>pH<sub>int</sub>=7.5<br>Temp=20°C |
| HDX time course (s) | 3, 100, 3000 at 20°C | 3, 100, 3000 at 20°C | 3, 100, 3000 at 20°C | 3, 100, 3000 at 20°C | 3, 100, 3000 at 20°C | 3, 100, 3000 at 20°C | 3, 100, 3000 at 20°C |
| HDX controls | N/A | N/A | N/A | N/A | N/A | N/A | N/A |
| Back-exchange | Corrected based on %D.O | Corrected based on %D.O | Corrected based on %D.O | Corrected based on %D.O | Corrected based on %D.O | Corrected based on %D.O | Corrected based on %D.O |
| Number of peptides | 16 | 26 | 21 | 16 | 29 | 26 | 19 |
| Sequence coverage | 84.1 | 88.6 | 86.4 | 60.0 | 91.3 | 82.4 | 87.9 |
| Average peptide /redundancy | Length=11.5<br>Redundancy=1.3 | Length=12.8<br>Redundancy=2.4 | Length=12.3<br>Redundancy=1.8 | Length=12<br>Redundancy=1.1 | Length=12.9<br>Redundancy=1.7 | Length=13.8<br>Redundancy=1.9 | Length=10.7<br>Redundancy=1.2 |
| Replicates | 3 | 3 | 3 | 3 | 3 | 3 | 3 |
| Repeatability | Average StDev=0.7% | Average StDev=0.6% | Average StDev=0.9% | Average StDev=0.6% | Average StDev=0.8% | Average StDev=0.9% | Average StDev=1% |
| Significant differences in HDX | >5% and >0.5 Da and unpaired t-test ≤0.01 | >5% and >0.5 Da and unpaired t-test ≤0.01 | >5% and >0.5 Da and unpaired t-test ≤0.01 | >5% and >0.5 Da and unpaired t-test ≤0.01 | >5% and >0.5 Da and unpaired t-test ≤0.01 | >5% and >0.5 Da and unpaired t-test ≤0.01 | >5% and >0.5 Da and unpaired t-test ≤0.01 |
| Data set | TRAPP C8 | TRAPP C9 | TRAPP C10 | TRAPP C11 | TRAPP C12 | TRAPP C13 |  |
| HDX reaction details | %D.O=62%<br>pH<sub>int</sub>=7.5<br>Temp=20°C | %D.O=62%<br>pH<sub>int</sub>=7.5<br>Temp=20°C | %D.O=62%<br>pH<sub>int</sub>=7.5<br>Temp=20°C | %D.O=62%<br>pH<sub>int</sub>=7.5<br>Temp=20°C | %D.O=62%<br>pH<sub>int</sub>=7.5<br>Temp=20°C | %D.O=62%<br>pH<sub>int</sub>=7.5<br>Temp=20°C |  |
| HDX time course (seconds) | 3, 100, 3000 at 20°C | 3, 100, 3000 at 20°C | 3, 100, 3000 at 20°C | 3, 100, 3000 at 20°C | 3, 100, 3000 at 20°C | 3, 100, 3000 at 20°C |  |
| HDX controls | N/A | N/A | N/A | N/A | N/A | N/A |  |
| Back-exchange | Corrected based on %D.O | Corrected based on %D.O | Corrected based on %D.O | Corrected based on %D.O | Corrected based on %D.O | Corrected based on %D.O |  |
| Number of peptides | 142 | 192 | 190 | 132 | 62 | 49 |  |
| Sequence coverage | 82.3 | 95.9 | 85 | 87.3 | 89.1 | 78.2 |  |
| Average peptide /redundancy | Length=15<br>Redundancy=1.5 | Length=13<br>Redundancy=2.2 | Length=13.6<br>Redundancy=2.0 | Length=12<br>Redundancy=1.4 | Length=14.4<br>Redundancy=2.0 | Length=10.8<br>Redundancy=1.3 |  |
| Replicates | 3 | 3 | 3 | 3 | 3 | 3 |  |
| Repeatability | Average StDev=1% | Average StDev=0.9% | Average StDev=0.8% | Average StDev=0.9% | Average StDev=0.8% | Average StDev=1% |  |
| Significant differences in HDX | >5% and >0.5 Da and unpaired t-test ≤0.01 | >5% and >0.5 Da and unpaired t-test ≤0.01 | >5% and >0.5 Da and unpaired t-test ≤0.01 | >5% and >0.5 Da and unpaired t-test ≤0.01 | >5% and >0.5 Da and unpaired t-test ≤0.01 | >5% and >0.5 Da and unpaired t-test ≤0.01 |  |

## References

[1] J. Cherfils, M. Zeghouf, Regulation of Small GTPases by GEFs, GAPs, and GDIs, Physiological Reviews. 93 (2013) 269–309. https://doi.org/10.1152/physrev.00003.2012.

[2] M.P. Müller, R.S. Goody, Molecular control of Rab activity by GEFs, GAPs and GDI, Small GTPases. 9 (2018) 5–21. https://doi.org/10.1080/21541248.2016.1276999.

[3] P. Novick, Regulation of membrane traffic by Rab GEF and GAP cascades, Small GTPases. 7 (2016) 252–256. https://doi.org/10.1080/21541248.2016.1213781.

[4] J. Barrowman, D. Bhandari, K. Reinisch, S. Ferro-Novick, TRAPP complexes in membrane traffic: convergence through a common Rab, Nature Reviews Molecular Cell Biology. 11 (2010) 759–763. https://doi.org/10.1038/nrm2999.

[5] S. Brunet, M. Sacher, In Sickness and in Health: The Role of TRAPP and Associated Proteins in Disease, Traffic. 15 (2014) 803–818. https://doi.org/10.1111/tra.12183.

[6] J.J. Kim, Z. Lipatova, N. Segev, TRAPP Complexes in Secretion and Autophagy, Front. Cell Dev. Biol. 4 (2016). https://doi.org/10.3389/fcell.2016.00020.

[7] Z. Lipatova, N. Segev, Ypt/Rab GTPases and their TRAPP GEFs at the Golgi, FEBS Letters. 593 (2019) 2488–2500. https://doi.org/10.1002/1873-3468.13574.

[8] M. Sacher, Y.-G. Kim, A. Lavie, B.-H. Oh, N. Segev, The TRAPP Complex: Insights into its Architecture and Function, Traffic. 9 (2008) 2032–2042. https://doi.org/10.1111/j.1600-0854.2008.00833.x.

[9] M. Sacher, N. Shahrzad, H. Kamel, M.P. Milev, TRAPPopathies: An emerging set of disorders linked to variations in the genes encoding transport protein particle (TRAPP)-associated proteins, Traffic. 20 (2019) 5–26. https://doi.org/10.1111/tra.12615.

[10] M.L. Jenkins, N.J. Harris, U. Dalwadi, K.D. Fleming, D.S. Ziemianowicz, A. Rafiei, E.M. Martin, D.C. Schriemer, C.K. Yip, J.E. Burke, The substrate specificity of the human TRAPPII complex’s Rab-guanine nucleotide exchange factor activity, Communications Biology. 3 (2020) 1–12. https://doi.org/10.1038/s42003-020-01459-2.

[11] F. Riedel, A. Galindo, N. Muschalik, S. Munro, The two TRAPP complexes of metazoans have distinct roles and act on different Rab GTPases, Journal of Cell Biology. 217 (2017) 601–617. https://doi.org/10.1083/jcb.201705068.

[12] L.L. Thomas, J.C. Fromme, GTPase cross talk regulates TRAPPII activation of Rab11 homologues during vesicle biogenesis, Journal of Cell Biology. 215 (2016) 499–513. https://doi.org/10.1083/jcb.201608123.

[13] N. Morozova, Y. Liang, A.A. Tokarev, S.H. Chen, R. Cox, J. Andrejic, Z. Lipatova, V.A. Sciorra, S.D. Emr, N. Segev, TRAPPII subunits are required for the specificity switch of a Ypt–Rab GEF, Nature Cell Biology. 8 (2006) 1263–1269. https://doi.org/10.1038/ncb1489.

[14] L.L. Thomas, A.M.N. Joiner, J.C. Fromme, The TRAPPIII complex activates the GTPase Ypt1 (Rab1) in the secretory pathway, Journal of Cell Biology. 217 (2017) 283–298. https://doi.org/10.1083/jcb.201705214.

[15] M.A. Lynch-Day, D. Bhandari, S. Menon, J. Huang, H. Cai, C.R. Bartholomew, J.H. Brumell, S. Ferro-Novick, D.J. Klionsky, Trs85 directs a Ypt1 GEF, TRAPPIII, to the phagophore to promote autophagy, PNAS. 107 (2010) 7811–7816. https://doi.org/10.1073/pnas.1000063107.

[16] K. Meiling-Wesse, U.D. Epple, R. Krick, H. Barth, A. Appelles, C. Voss, E.-L. Eskelinen, M. Thumm, Trs85 (Gsg1), a Component of the TRAPP Complexes, Is Required for the Organization of the Preautophagosomal Structure during Selective Autophagy via the Cvt Pathway*, Journal of Biological Chemistry. 280 (2005) 33669–33678. https://doi.org/10.1074/jbc.M501701200.

[17] C. Li, Z. Wei, Y. Fan, W. Huang, Y. Su, H. Li, Z. Dong, M. Fukuda, M. Khater, G. Wu, The GTPase Rab43 Controls the Anterograde ER-Golgi Trafficking and Sorting of GPCRs, Cell Rep. 21 (2017) 1089–1101. https://doi.org/10.1016/j.celrep.2017.10.011.

[18] Y.-G. Kim, S. Raunser, C. Munger, J. Wagner, Y.-L. Song, M. Cygler, T. Walz, B.-H. Oh, M. Sacher, The Architecture of the Multisubunit TRAPP I Complex Suggests a Model for Vesicle Tethering, Cell. 127 (2006) 817–830. https://doi.org/10.1016/j.cell.2006.09.029.

[19] M.C. Bassik, M. Kampmann, R.J. Lebbink, S. Wang, M.Y. Hein, I. Poser, J. Weibezahn, M.A. Horlbeck, S. Chen, M. Mann, A.A. Hyman, E.M. LeProust, M.T. McManus, J.S. Weissman, A Systematic Mammalian Genetic Interaction Map Reveals Pathways Underlying Ricin Susceptibility, Cell. 152 (2013) 909–922. https://doi.org/10.1016/j.cell.2013.01.030.

[20] P.J. Scrivens, B. Noueihed, N. Shahrzad, S. Hul, S. Brunet, M. Sacher, C4orf41 and TTC-15 are mammalian TRAPP components with a role at an early stage in ER-to-Golgi trafficking, MBoC. 22 (2011) 2083–2093. https://doi.org/10.1091/mbc.e10-11-0873.

[21] C.A. Lamb, S. Nühlen, D. Judith, D. Frith, A.P. Snijders, C. Behrends, S.A. Tooze, TBC1D14 regulates autophagy via the TRAPP complex and ATG9 traffic, The EMBO Journal. 35 (2016) 281–301. https://doi.org/10.15252/embj.201592695.

[22] Y. Cai, H.F. Chin, D. Lazarova, S. Menon, C. Fu, H. Cai, A. Sclafani, D.W. Rodgers, E.M. De La Cruz, S. Ferro-Novick, K.M. Reinisch, The Structural Basis for Activation of the Rab Ypt1p by the TRAPP Membrane-Tethering Complexes, Cell. 133 (2008) 1202–1213. https://doi.org/10.1016/j.cell.2008.04.049.

[23] A.M.N. Joiner, B.P. Phillips, K. Yugandhar, E.J. Sanford, M.B. Smolka, H. Yu, E.A. Miller, J.C. Fromme, Structure and mechanism of TRAPPIII-mediated Rab1 activation, BioRxiv. (2020) 2020.10.08.332312. https://doi.org/10.1101/2020.10.08.332312.

[24] A. Galindo, V.J. Planelles-Herrero, G. Degliesposti, S. Munro, A cryo-EM structure of metazoan TRAPPIII, the multisubunit complex that activates the GTPase Rab1, BioRxiv. (2020) 2020.12.17.423307. https://doi.org/10.1101/2020.12.17.423307.

[25] F. Weissmann, G. Petzold, R. VanderLinden, P.J.H. in ’t Veld, N.G. Brown, F. Lampert, S. Westermann, H. Stark, B.A. Schulman, J.-M. Peters, biGBac enables rapid gene assembly for the expression of large multisubunit protein complexes, PNAS. 113 (2016) E2564–E2569. https://doi.org/10.1073/pnas.1604935113.

[26] L.A. Kelley, S. Mezulis, C.M. Yates, M.N. Wass, M.J.E. Sternberg, The Phyre2 web portal for protein modeling, prediction and analysis, Nature Protocols. 10 (2015) 845–858. https://doi.org/10.1038/nprot.2015.053.

[27] L.L. Thomas, S.A. van der Vegt, J.C. Fromme, A Steric Gating Mechanism Dictates the Substrate Specificity of a Rab-GEF, Developmental Cell. 48 (2019) 100–114.e9. https://doi.org/10.1016/j.devcel.2018.11.013.

[28] T.H. Klöpper, N. Kienle, D. Fasshauer, S. Munro, Untangling the evolution of Rab G proteins: implications of a comprehensive genomic analysis, BMC Biol. 10 (2012) 71. https://doi.org/10.1186/1741-7007-10-71.

[29] W. Cheng, K. Yin, D. Lu, B. Li, D. Zhu, Y. Chen, H. Zhang, S. Xu, J. Chai, L. Gu, Structural insights into a unique Legionella pneumophila effector LidA recognizing both GDP and GTP bound Rab1 in their active state, PLoS Pathog. 8 (2012) e1002528. https://doi.org/10.1371/journal.ppat.1002528.

[30] M.L. Jenkins, J.P. Margaria, J.T.B. Stariha, R.M. Hoffmann, J.A. McPhail, D.J. Hamelin, M.J. Boulanger, E. Hirsch, J.E. Burke, Structural determinants of Rab11 activation by the guanine nucleotide exchange factor SH3BP5, Nat Commun. 9 (2018) 3772. https://doi.org/10.1038/s41467-018-06196-z.

[31] D. Taussig, Z. Lipatova, N. Segev, Trs20 is Required for TRAPP III Complex Assembly at the PAS and its Function in Autophagy, Traffic. 15 (2014) 327–337. https://doi.org/10.1111/tra.12145.

[32] M. Pinar, E. Arias-Palomo, V. de los Ríos, H.N.A. Jr, M.A. Peñalva, Characterization of Aspergillus nidulans TRAPPs uncovers unprecedented similarities between fungi and metazoans and reveals the modular assembly of TRAPPII, PLOS Genetics. 15 (2019) e1008557. https://doi.org/10.1371/journal.pgen.1008557.

[33] D. Tan, Y. Cai, J. Wang, J. Zhang, S. Menon, H.-T. Chou, S. Ferro-Novick, K.M. Reinisch, T. Walz, The EM structure of the TRAPPIII complex leads to the identification of a requirement for COPII vesicles on the macroautophagy pathway, PNAS. 110 (2013) 19432–19437. https://doi.org/10.1073/pnas.1316356110.

[34] C.K. Yip, J. Berscheminski, T. Walz, Molecular architecture of the TRAPPII complex and implications for vesicle tethering, Nature Structural & Molecular Biology. 17 (2010) 1298–1304. https://doi.org/10.1038/nsmb.1914.

[35] S. Jones, C. Newman, F. Liu, N. Segev, The TRAPP Complex Is a Nucleotide Exchanger for Ypt1 and Ypt31/32, MBoC. 11 (2000) 4403–4411. https://doi.org/10.1091/mbc.11.12.4403.

[36] Z. Lipatova, N.V. Bergen, D. Stanga, M. Sacher, J. Christodoulou, N. Segev, TRAPPing a neurological disorder: from yeast to humans, Autophagy. 16 (2020) 965–966. https://doi.org/10.1080/15548627.2020.1736873.

[37] M. Zong, X. Wu, C.W.L. Chan, M.Y. Choi, H.C. Chan, J.A. Tanner, S. Yu, The Adaptor Function of TRAPPC2 in Mammalian TRAPPs Explains TRAPPC2-Associated SEDT and TRAPPC9-Associated Congenital Intellectual Disability, PLOS ONE. 6 (2011) e23350. https://doi.org/10.1371/journal.pone.0023350.

[38] S. Brunet, N. Shahrzad, D. Saint-Dic, H. Dutczak, M. Sacher, A trs20 Mutation That Mimics an SEDT-Causing Mutation Blocks Selective and Non-Selective Autophagy: A Model for TRAPP III Organization, Traffic. 14 (2013) 1091–1104. https://doi.org/10.1111/tra.12095.

[39] M.P. Milev, C. Graziano, D. Karall, W.F.E. Kuper, N. Al-Deri, D.M. Cordelli, T.B. Haack, K. Danhauser, A. Iuso, F. Palombo, T. Pippucci, H. Prokisch, D. Saint-Dic, M. Seri, D. Stanga, G. Cenacchi, K.L.I. van Gassen, J. Zschocke, C. Fauth, J.A. Mayr, M. Sacher, P.M. van Hasselt, Bi-allelic mutations in TRAPPC2L result in a neurodevelopmental disorder and have an impact on RAB11 in fibroblasts, Journal of Medical Genetics. 55 (2018) 753–764. https://doi.org/10.1136/jmedgenet-2018-105441.

[40] S. Duerinckx, M. Meuwissen, C. Perazzolo, L. Desmyter, I. Pirson, M. Abramowicz, Phenotypes in siblings with homozygous mutations of TRAPPC9 and/or MCPH1 support a bifunctional model of MCPH1, Molecular Genetics & Genomic Medicine. 6 (2018) 660–665. https://doi.org/10.1002/mgg3.400.

[41] D.B. Fee, M. Harmelink, P. Monrad, E. Pyzik, Siblings With Mutations in TRAPPC11 Presenting With Limb-Girdle Muscular Dystrophy 2S, Journal of Clinical Neuromuscular Disease. 19 (2017) 27–30. https://doi.org/10.1097/CND.0000000000000173.

[42] A.A. Larson, P.R. Baker, M.P. Milev, C.A. Press, R.J. Sokol, M.O. Cox, J.K. Lekostaj, A.A. Stence, A.D. Bossler, J.M. Mueller, K. Prematilake, T.F. Tadjo, C.A. Williams, M. Sacher, S.A. Moore, TRAPPC11 and GOSR2 mutations associate with hypoglycosylation of α-dystroglycan and muscular dystrophy, Skeletal Muscle. 8 (2018) 17. https://doi.org/10.1186/s13395-018-0163-0.

[43] F. Weissmann, G. Petzold, R. VanderLinden, P.J. Huis in ’t Veld, N.G. Brown, F. Lampert, S. Westermann, H. Stark, B.A. Schulman, J.-M. Peters, biGBac enables rapid gene assembly for the expression of large multisubunit protein complexes, Proceedings of the National Academy of Sciences. 113 (2016) E2564–E2569. https://doi.org/10.1073/pnas.1604935113.

[44] A. Delprato, E. Merithew, D.G. Lambright, Structure, exchange determinants, and family-wide rab specificity of the tandem helical bundle and Vps9 domains of Rabex-5, Cell. 118 (2004). https://doi.org/10.1016/j.cell.2004.08.009.

[45] J.T.B. Stariha, R.M. Hoffmann, D.J. Hamelin, J.E. Burke, Probing Protein-Membrane Interactions and Dynamics Using Hydrogen-Deuterium Exchange Mass Spectrometry (HDX-MS), Methods Mol Biol. 2263 (2021) 465–485. https://doi.org/10.1007/978-1-0716-1197-5_22.

[46] J.M. Dobbs, M.L. Jenkins, J.E. Burke, Escherichia coli and Sf9 Contaminant Databases to Increase Efficiency of Tandem Mass Spectrometry Peptide Identification in Structural Mass Spectrometry Experiments, J Am Soc Mass Spectrom. 31 (2020) 2202–2209. https://doi.org/10.1021/jasms.0c00283.

[47] G.R. Masson, J.E. Burke, N.G. Ahn, G.S. Anand, C. Borchers, S. Brier, G.M. Bou-Assaf, J.R. Engen, S.W. Englander, J. Faber, R. Garlish, P.R. Griffin, M.L. Gross, M. Guttman, Y. Hamuro, A.J.R. Heck, D. Houde, R.E. Iacob, T.J.D. Jørgensen, I.A. Kaltashov, J.P. Klinman, L. Konermann, P. Man, L. Mayne, B.D. Pascal, D. Reichmann, M. Skehel, J. Snijder, T.S. Strutzenberg, E.S. Underbakke, C. Wagner, T.E. Wales, B.T. Walters, D.D. Weis, D.J. Wilson, P.L. Wintrode, Z. Zhang, J. Zheng, D.C. Schriemer, K.D. Rand, Recommendations for performing, interpreting and reporting hydrogen deuterium exchange mass spectrometry (HDX-MS) experiments, Nature Methods. 16 (2019) 595–602. https://doi.org/10.1038/s41592-019-0459-y.

[48] Y. Perez-Riverol, A. Csordas, J. Bai, M. Bernal-Llinares, S. Hewapathirana, D.J. Kundu, A. Inuganti, J. Griss, G. Mayer, M. Eisenacher, E. Pérez, J. Uszkoreit, J. Pfeuffer, T. Sachsenberg, Ş. Yılmaz, S. Tiwary, J. Cox, E. Audain, M. Walzer, A.F. Jarnuczak, T. Ternent, A. Brazma, J.A. Vizcaíno, The PRIDE database and related tools and resources in 2019: improving support for quantification data, Nucleic Acids Research. 47 (2019) D442–D450. https://doi.org/10.1093/nar/gky1106.

[49] A. Rohou, N. Grigorieff, CTFFIND4: Fast and accurate defocus estimation from electron micrographs, Journal of Structural Biology. 192 (2015) 216–221. https://doi.org/10.1016/j.jsb.2015.08.008.

[50] New tools for automated high-resolution cryo-EM structure determination in RELION-3 | eLife, (n.d.). https://elifesciences.org/articles/42166 (accessed May 14, 2021).

[51] A. Punjani, J.L. Rubinstein, D.J. Fleet, M.A. Brubaker, cryoSPARC: algorithms for rapid unsupervised cryo-EM structure determination, Nature Methods. 14 (2017) 290–296. https://doi.org/10.1038/nmeth.4169.

## References

[1] X. Robert, P. Gouet, Deciphering key features in protein structures with the new ENDscript server, Nucleic Acids Res. 42 (2014) W320–324. https://doi.org/10.1093/nar/gku316.

[2] M.L. Jenkins, N.J. Harris, U. Dalwadi, K.D. Fleming, D.S. Ziemianowicz, A. Rafiei, E.M. Martin, D.C. Schriemer, C.K. Yip, J.E. Burke, The substrate specificity of the human TRAPPII complex’s Rab-guanine nucleotide exchange factor activity, Communications Biology. 3 (2020) 1–12. https://doi.org/10.1038/s42003-020-01459-2.

